# Single-cell time-series mapping of cell fate trajectories reveals an expanded developmental potential for human PSC-derived distal lung progenitors

**DOI:** 10.1101/782896

**Authors:** Killian Hurley, Jun Ding, Carlos Villacorta-Martin, Michael J. Herriges, Anjali Jacob, Marall Vedaie, Konstantinos D. Alysandratos, Yuliang L. Sun, Chieh Lin, Rhiannon B. Werder, Andrew A. Wilson, Aditya Mithal, Gustavo Mostoslavsky, Ignacio S. Caballero, Susan H. Guttentag, Farida Ahangari, Naftali Kaminski, Alejo Rodriguez-Fraticelli, Fernando Camargo, Ziv Bar-Joseph, Darrell N Kotton

**Affiliations:** Center for Regenerative Medicine of Boston University and Boston Medical Center, Boston, MA 02118, USA; The Pulmonary Center and Department of Medicine, Boston University School of Medicine, Boston, MA 02118, USA; Department of Medicine, Royal College of Surgeons in Ireland, Education and Research Centre, Beaumont Hospital, Dublin, Ireland; Tissue Engineering Research Group, Royal College of Surgeons in Ireland, Ireland; Computational Biology Department, School of Computer Science, Carnegie Mellon University, Pittsburgh, PA 15213, USA; Machine Learning Department, School of Computer Science, Carnegie Mellon University, Pittsburgh, PA 15217, USA; Department of Pediatrics; Monroe Carell Jr. Children’s Hospital, Vanderbilt University, Nashville, TN 37232, USA; Pulmonary, Critical care and Sleep Medicine, Yale University School of Medicine, New Haven, CT 16520, USA; Stem Cell Program, Boston Children’s Hospital, Boston, MA 02115, USA; Department of Stem Cell and Regenerative Biology, Harvard University, Cambridge, MA 02138, USA; Harvard Stem Cell Institute, Boston, MA 02115, USA

## Abstract

Alveolar epithelial type 2 cells (AEC2s) are the facultative progenitors responsible for maintaining lung alveoli throughout life, yet are difficult to access from patients for biomedical research or lung regeneration applications. Here we engineer AEC2s from human induced pluripotent stem cells (iPSCs) in vitro and use single cell RNA sequencing (scRNA-seq) to profile the detailed kinetics of their differentiation over time. We focus on both the desired target cells as well as those that appear to diverge to alternative endodermal fates. By combining scRNA-seq with lentiviral barcoding to trace differentiating clones, we reveal the bifurcating cell fate trajectories followed as primordial lung progenitors differentiate into mature AEC2s. We define the global transcriptomic signatures of primary developing human AEC2s from fetal through adult stages in order to identify the subset of in vitro differentiating cells that appear to recapitulate the path of in vivo development. In addition, we develop computational methods based on Continuous State Hidden Markov Models (CSHMM) to identify the precise timing and type of signals, such as over-exuberant Wnt responses, that induce some early multipotent NKX2-1+ progenitors to lose lung fate as they clonally diverge into a variety of non-lung endodermal lineages. Finally, we find that this initial developmental plasticity is regulatable via Wnt modulation, and subsides over time, ultimately resulting in iPSC-derived AEC2s that exhibit a stable phenotype and nearly limitless self-renewal capacity in vitro. Our methods and computational approaches can be widely applied to study and control directed differentiation, producing an inexhaustible supply of mature lineages, exemplified here by the generation of AEC2s.

## Introduction

A central aim of developmental biology is to better understand the embryonic differentiation and maturation pathways that lead to functioning adult cells and tissues. Multistage, step-wise differentiation protocols applied to cultured human pluripotent stem cells (PSC) are designed to recapitulate these pathways in order to produce specific mature target cells. This approach allows the detailed in vitro study of the kinetics of human development at embryonic time points that are difficult to access in vivo, while also producing populations of cells for regenerative therapies and disease modelling. However, even the most optimized PSC differentiation protocols tend to yield a complex, heterogenous mix of cells of varying fates and maturation states, limiting the successful recapitulation of target cell identity or purity (Schwartzentruber et al., 2018; Wu et al., 2018). This hurdle makes it challenging to understand the molecular mechanisms underlying human in vivo differentiation and consequently leads to limited clinical relevance and utility for several PSC-derived lineages.

The study of human lung development exemplifies this challenge. Access to developing fetal primary cells as experimental controls is limited, while in vitro differentiation of PSCs must attempt to recapitulate at least 20 weeks of gestational time that elapses from the moment of in vivo lung epithelial endodermal specification (approximately 4 weeks) until maturation of the earliest distal lung alveolar epithelial cells that exhibit surfactant producing organelles (24 weeks). We and others have published in vitro PSC directed differentiation protocols which reduce the duration of this analogous in vivo developmental window to 2 weeks in vitro as PSC-derived lung endodermal precursors (NKX2-1+ primordial lung progenitors) are differentiated in culture into lung alveolar epithelial type 2 cells (iAEC2s), the life-long facultative progenitors of the alveolar epithelium (Gotoh et al., 2014; Huang et al., 2014; Jacob et al., 2017). We have further shown that despite sorting to purity for the earliest known lung progenitors, identified by NKX2-1 expression, during directed differentiation (Hawkins et al., 2017) the resulting cells are plastic, transcriptomically heterogenous, and tend to drift over time into a variety of both lung and non-lung molecular phenotypes (McCauley et al., 2018). Importantly, this finding of fate heterogeneity mimics a variety of *in vivo* mouse developmental models or human lung cancer settings, which also document the emergence of ectopic endodermal programs in lung epithelial cells if signalling pathways, such as Wnt, or gene regulatory networks, such as those downstream of NKX2-1, are perturbed during key stages of fetal or adult life (Okubo and Hogan, 2004; Snyder et al., 2013; Tata et al., 2018). Given these results, detailed mapping of the developmental path or paths that progenitor cells take during differentiation to their end state or fate both in vivo as well as in PSC-derived systems has now become a primary objective of the field. Similar challenges in obtaining pure cell populations from PSC differentiation protocols were also recently observed for other tissue types such as the renal epithelium (Holtzinger et al., 2015; Schwartzentruber et al., 2018; Wu et al., 2018).

While single cell RNA sequencing (scRNA-seq) can provide a detailed picture of cell states, distinguishing between immature and fully differentiated PSC-derived cells, this technique alone loses information about spatial and temporal factors and can only imply cell parent-progeny relationships in the absence of a lineage tracing strategy (Weinreb et al., 2018b). Several methods have been developed for inferring single cell trajectories (Trapnell et al., 2014); however, these usually rely on dimensionality reduction which make it hard to infer the regulatory process that controls the branching of various cell fates (Ding et al., 2018).

To address these issues here we present a general strategy for modelling such trajectories that can be used to better understand and improve differentiation protocols. We first employ bulk RNA sequencing of primary developing fetal and adult lung cells in order to map in vivo developmental maturation over time, establishing benchmark datasets and verifying key signalling pathways associated with maturation of the differentiated cells. Next, using a computational algorithm to interrogate the expression kinetics of a subset of genes profiled first at high resolution in differentiating PSCs, we select a set of optimal time points for global transcriptomic profiling and for these perform scRNA-seq time series analyses of PSC-derived cells. We use a novel computational method based on Continuous State Hidden Markov Models (CSHMM) to construct developmental trajectories and to identify the regulators and pathways involved in controlling the process. We then use the computational model to predict both the *type* and *timing* of potential interventions which can be used to increase the fraction of cells branching to the desired fate, and we combine lentiviral barcoding with scRNA-seq to validate the parent-progeny lineage relationships and fate bifurcations predicted by our model. The outcome of these studies is a markedly improved understanding of the kinetics, fate trajectories, and cellular plasticity associated with in vitro human PSC directed differentiation, exemplified here by the derivation of lung alveolar epithelial cells from their developmental endodermal precursors.

## Results

### Transcriptomic profiles of primary human developing lung alveolar epithelium

We first sought to identify the transcriptional kinetics of maturation that characterize *in vivo* development of human AEC2s. We performed bulk RNA sequencing (RNA-seq) of distal human fetal and adult lung alveolar epithelium at 3 key developmental time points (Figure 1A). Using our previously published methods (Gonzales et al., 2002; Jacob et al., 2017; Wade et al., 2006), we purified alveolar epithelial cells from human fetal lungs (HFL) at 16-17.5 weeks of gestation (n=3; hereafter Early HFL), 20-21 weeks of gestation (n=4; hereafter Late HFL), and postnatally from adult lungs (n=3). These three time points represent, respectively: a) early canalicular-staged cells composed of distal lung bud tip epithelial cells that are thought to have already initiated their alveolar programs; (Miller et al., 2018; Nikolic et al., 2017; Nikolic et al., 2018); b) more differentiated alveolar cells at a later canalicular stage but just prior to the emergence of lamellar bodies (Nikolic et al., 2017), and c) fully mature adult AEC2s, sorted based on HTII-280 expression (Jacob et al., 2017). To enable comparison of these samples to an earlier staged NKX2-1+ lung endodermal progenitor, we also profiled in vitro PSC-derived “primordial lung endodermal progenitors” (PLP) (n=3), sorted using previously published surface markers (CD47hi/CD26lo) (Hawkins et al., 2017). At an FDR<0.05 (empirical Bayes ANOVA test) we identified 15,137 differentially expressed genes across the 13 samples, in line with recent studies that found that a large portion of the transcriptome is differentially expressed in early development (Junyue Cao et al., 2019).

**Figure 1.**
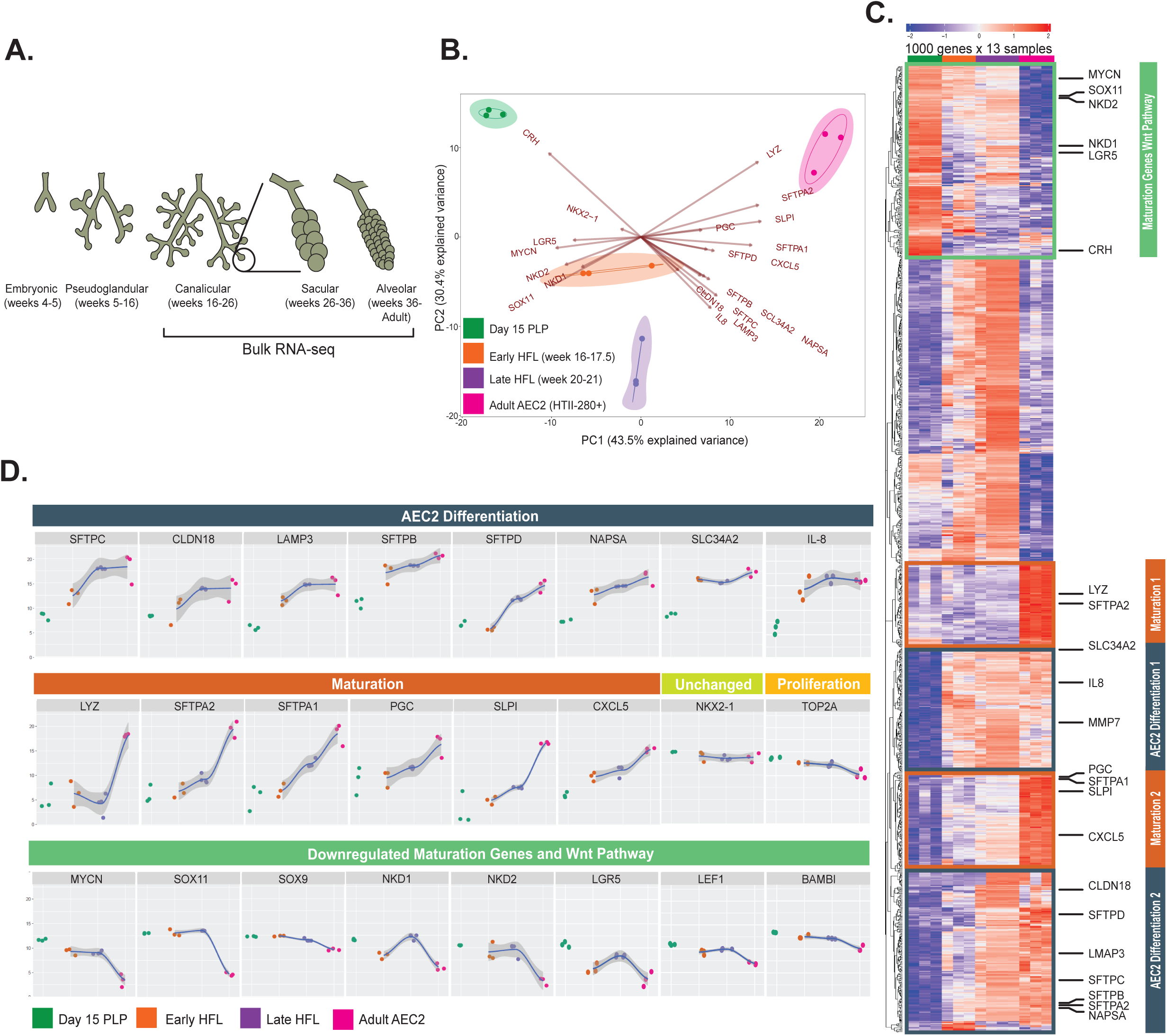
Global transcriptomic time series reveals the kinetics of developing primary human alveolar epithelial type 2 cells. (A) The 5 stages of human lung development highlighting the samples obtained for bulk RNA sequencing. (B) Principal component analysis (PCA) of gene expression (18,147 detected genes) across all 13 samples including primordial lung progenitors (PLP) derived from pluripotent stem cells at day 15 of differentiation, primary early human fetal lung (HFL; 16-17.5 weeks gestation), late HFL (20-21 weeks gestation), and adult alveolar epithelial type 2 cells (AEC2) sorted on the antibody HTII-280. The loadings of highly variable genes associated with differentiation and maturation of AEC2 in panel C, are overlayed on the PCA plot in B to indicate their weight on PC1 and PC2. Arrow tips denote the correlation coefficient of the respective gene with each principal component. (C) Heatmap showing unsupervised hierarchical clustering of the top 1,000 most variable genes across all samples. Differentiation and maturation markers are highlighted on the right. (D) Smoothed regressions of time series samples indicating normalized gene expression values (from panel C) for 8 selected genes associated with differentiation of AEC2, 6 genes associated with maturation of AEC2 and 8 selected downregulated AEC2 maturation and Wnt pathway genes. See also Table S1

To focus on a smaller number of genes, we selected the 1000 genes with highest variance in expression across all 13 samples in order to identify transcripts most associated with early human alveolar differentiation vs maturation (Figure 1B, C). Hierarchical clustering of these genes identified candidate “Differentiation” clusters which included a cluster that varied early in alveolar development (weeks 16-21) and a “Maturation” cluster enriched preferentially in adult AEC2s (figure 1C). We plotted the expression kinetics of cluster genes to select 8 markers of early distal alveolar differentiation that are expressed during fetal canalicular stages prior to full AEC2 maturation (“Differentiation” gene set: SFTPB, SFTPC, SFTPD, CLDN18, LAMP3, SLC34A2, IL8, and NAPSA; Figure 1D). Next, we identified a 6 gene “maturation” marker set, LYZ, SFTPA1, SFTPA2, PGC, CXCL5, and SLPI, which was preferentially expressed in adult AEC2s. We further identified genes downregulated in adult AEC2s (MYCN, SOX11, and the Wnt target genes, NKD1, NKD2, and LGR5 Figure 1D). Based on prior literature (Frank et al., 2016; Hogan et al., 2014; Jacob et al., 2017), and significant differential expression between primary fetal and adult AEC2s, we also selected downregulation of Wnt targets LEF1 and BAMBI and the transcription factor SOX9 as additional stage-dependent maturation markers of AEC2s, although SOX9 variance was not in the top 1000 varying genes overall. Taken together our profiles indicate the early development of distal human lung epithelium is characterized by the expression of a subset of surfactant- and lamellar body-associated genes, some of which increase non-linearly over time (Figure 1C and D), followed by later expression of genes associated with AEC2 maturation, including expression of the full complement of surfactant proteins and additional markers that others have observed in adult AEC2s (Desai et al., 2014; Treutlein et al., 2014)(Guo et al., 2019). Conversely, maturation of AEC2s is associated with decreasing Wnt signalling consistent with prior findings in vivo in mice (Frank et al., 2016) as well is in vitro in AEC2s (iAEC2) (Jacob et al., 2017). In contrast, the canonical transcription factor required for lung epithelial development, NKX2-1, maintains its expression over time (Figure 1D) in developing iAEC2s, supporting its utility as a marker expressed throughout the lifetime of AEC2s.

### Selecting the most appropriate time points to profile in a scRNA-seq analysis

Using the markers identified from profiling the transcriptomic kinetics of primary alveolar epithelial differentiation, we next sought to determine whether the differentiation trajectories of PSC-derived cells truly recapitulate human lung developmental kinetics. We employed our recently published protocol (Jacob et al., 2017), differentiating purified PSC-derived primordial NKX2-1+ lung progenitors over a 2-week period into iAEC2s. This prolonged time needed to differentiate human iAEC2s presents a substantial problem in selecting the number of developmental time points to exhaustively profile using costly methods such as scRNA-seq. The question of when and how often to sample is particularly challenging in developmental in vitro models, as molecular changes are likely non-linear, so simple linear sampling may fail to identify significant gene fluxes (Li et al., 2013). To address this issue, we adapted an algorithm we previously developed for the task of selecting optimal time points to profile in scRNA-seq studies using bulk data containing gene subsets. Our algorithm, Time Point Selection (TPS) (Kleyman et al., 2017), profiles a small set of selected genes sampled at a high rate. These are represented using splines and a combinatorial search is applied to select a subset of suitable points so that combined, selected points provide enough information to reconstruct the values for all genes across all time points (including those not selected, Figure 1C). The final number of points to be used can be determined as a function of the reconstructed error. To use TPS we profiled 80 relevant differentiation and maturation genes in PSC-derived differentiation to iAEC2s, every 2 days, over a 16-day period by NanoString (Figure 2B). Genes were selected from our in vivo analysis discussed above and from prior endodermal profiling (Hawkins et al., 2017) (Table S1 and Figure S1). For these studies, we utilized a non-diseased iPSC line (BU3 NGST) that we have engineered to carry knock in fluorescent reporters (NKX2-1^GFP^; SFTPC^tdTomato^) allowing real time monitoring of cells as their cell states proceed from initial lung specification (NKX2-1^GFP+^) through their differentiation into NKX2-1^GFP+/^SFTPC^tdTomato+^ iAEC2s (Jacob et al., 2017). We separated lung from non-lung cells at the primordial progenitor stage (based on GFP+ vs – sorting; Figure 2A) and profiled the outgrowth of the GFP+ vs GFP-populations without further cell sorting (Figure 2C and Figure S1).

**Figure 2.**
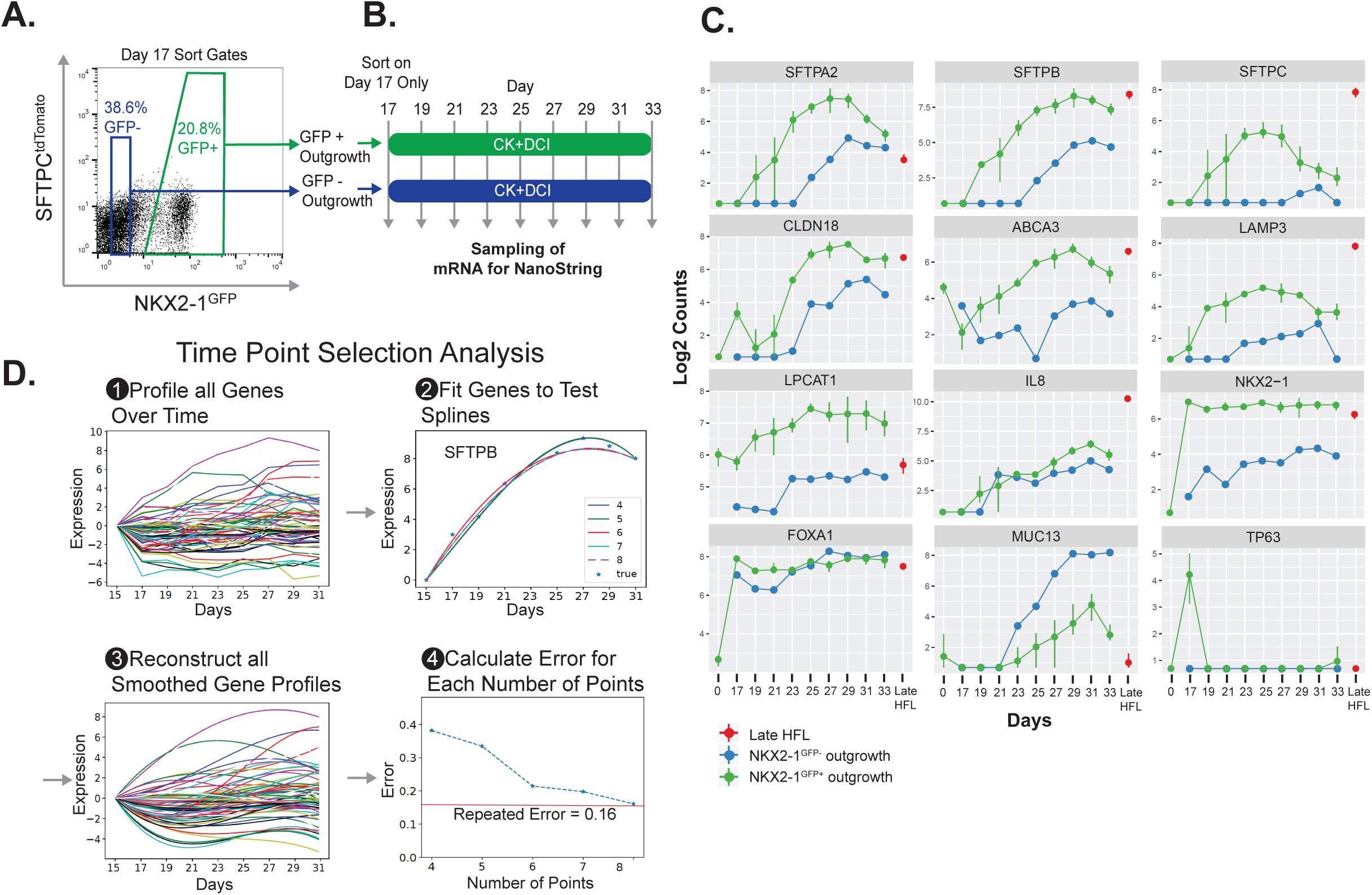
Time point selection (TPS) analysis of lung differentiation by NanoString predicts optimal time points for global transcriptomic scRNS-seq profiling. (A) Representative sort gates of iPSCs (BU3 NGST) sorted on NKX2-1^GFP^+ vs. NKX2-1^GFP^-on day 17 of differentiation for replating and time series NanoString analyses of their resulting outgrowths over time. (B) Schematic of differentiation protocol after flow cytometry sorting at day 17, showing time points for sampling for NanoString mRNA profiling of sorted NKX2-1^GFP^+ or sorted NKX2-1^GFP^-outgrowths. (C) Average and range of expression over time for selected genes associated with differentiation and maturation of AEC2 derived from PSCs (0 to 33 days of differentiation, n=3 biological replicates). Red dot indicates late human fetal lung controls (HFL; 20-21 weeks gestation). (D) Method for choosing the appropriate time points of the single-cell experiment. 1) 66 genes were profiled at high frequency using bulk cultured samples 2) regression splines are fitted in order to 3) model the expression of each gene and 4) iteratively evaluate the effect of removing time-points on the overall error until an optimal is found.

For this small panel we observed that expression of surfactant-encoding and lamellar body-related genes, SFTPB, SFTPC, SFTPA2, ABCA3, CLDN18 and LAMP3, increased over time to a maximum at days 25-29 and deceased thereafter (Figure 2C), while NKX2-1 remained constant. As expected, lung epithelial markers were relatively depleted in the outgrowth of GFP negative controls, consistent with our prior reports that all PSC-derived human lung epithelial lineages derive via the gateway of an NKX2-1+ primordial progenitor stage (Hawkins et al., 2017; Jacob et al., 2017; McCauley et al., 2018; McCauley et al., 2017; Serra et al., 2017). TPS identified an inflection point when using the optimal 6 time points (days 15 17, 21, 25, 29 and 31). While error increased rapidly when using less than 6 points, 7 or more points did not significantly reduce reconstruction error. As seen in Figure 2D, for the optimal set of 6 points reconstruction error is close to repeat error suggesting accurate inference of non-profiled time points.

### A single cell map of PSC-derived distal lung differentiation implies fate trajectories

We next profiled the transcriptional trajectories of individual cell states over time by scRNA-seq performed at the time intervals selected by our TPS algorithm (Figure 3A). For each time point we profiled ∼4000 cells, following PSC-derived cells sorted on NKX2-1^GFP^ at the lung primordial progenitor stage of differentiation (day 15; Figure 3B) through day 31 of alveolar directed differentiation without further sorting. As a negative control we included a 7^th^ sample, the day 15 non lung population, isolated based on GFP exclusion (NKX2-1^GFP^ negative sorted). Flow cytometry monitoring of NKX2-1 and SFTPC locus activity at each of the 6 time points of alveolar differentiation (Figure 3B) indicated the expected emergence of SFTPC^tdTomato^ expression over time in some cells, peaking on day 29 of differentiation, but a loss of NKX2-1^GFP^ expression in other cells over time, predicting potential loss of lung cell fate in a subset of the population, consistent with our prior reports (McCauley et al., 2018).

**Figure 3.**
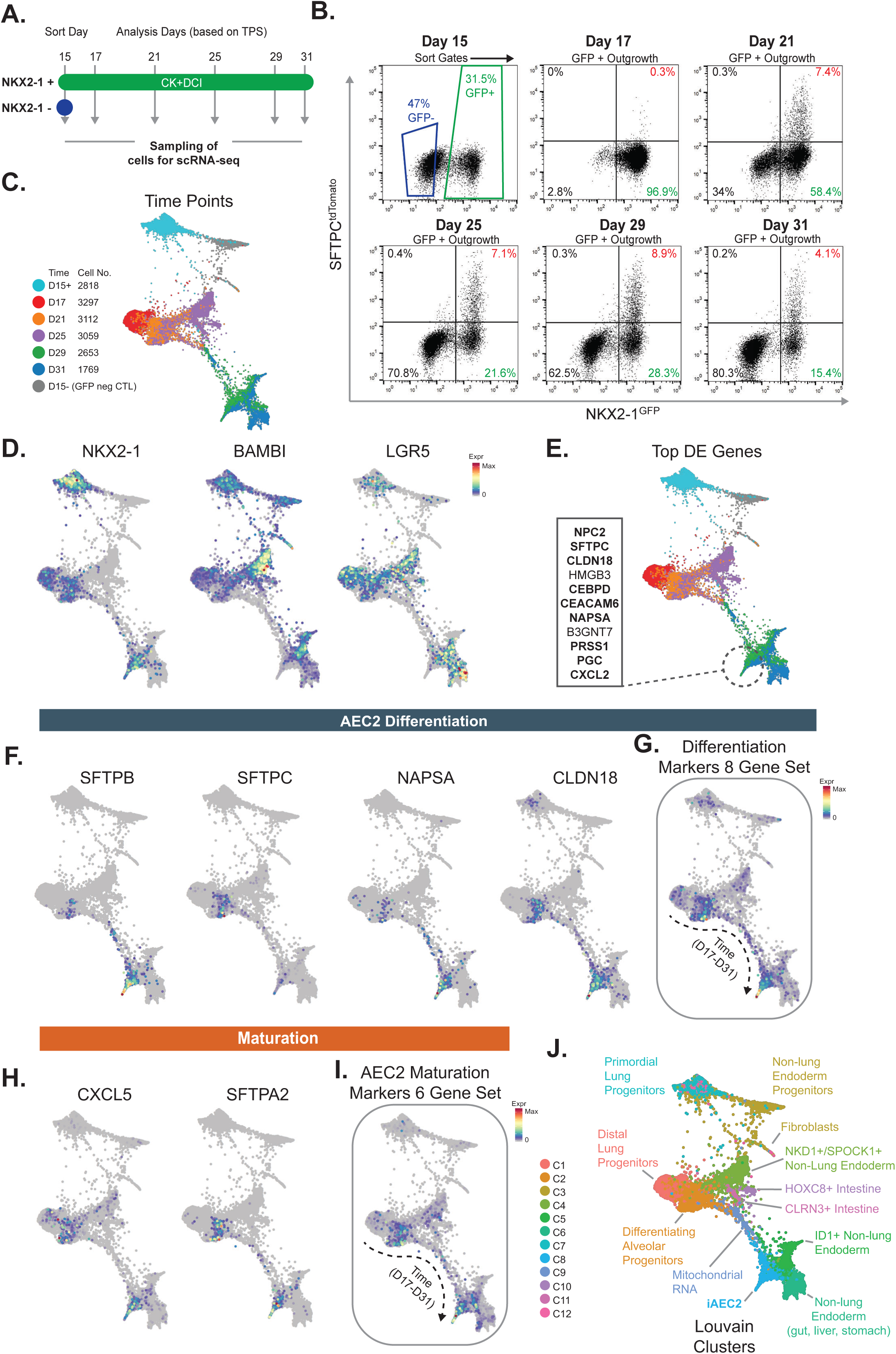
Time series single-cell transcriptomic analysis of AEC2 directed differentiation. (A) Schematic of experiment indicating time points for initial sorting of iPSC-derived primordial lung progenitors (day 15) and analysis of their outgrowths over time. (B) Flow cytometry sort gates and analyses based on expression of NKX2-1^GFP^ and SFTPC^tdTomato^ reporters at the time of harvest for each scRNA-seq analysis. (C) SPRING analysis of all cells across all 6 time points. “D15-“ represents the NKX2-1^GFP^ negative control shown in A, sorted on day 15 for comparison. All other time points indicate the outgrowth of GFP+ cells sorted on day 15 (D15+). (D) Normalized gene expression overlayed on SPRING plots for selected markers of retained lung fate (NKX2-1) vs BMP and Wnt signalling markers that are highly expressed in cells that appear to lose lung fate over time. (E) The top 11 transcripts upregulated in cells in the indicated gate compared to all other cells, bold font indicates a known marker of AEC2s. (F-I) Normalized expression levels for selected AEC2 marker genes as well as the composite set (G) of 8 AEC2 differentiation marker genes or 6 AEC2 maturation marker genes (H, I) from Figure 1. (J) Twelve cell clusters identified by Louvain clustering with identities assigned based on markers explained in the text or indicated in the panel. See also Figure 1 and Figure S2.

To visualize potential single cell fate trajectories in our model while preserving high-dimensional relationships we first utilized the SPRING algorithm (Weinreb et al., 2018a) to prepare force-directed layouts of k-nearest neighbour graphs for the entire differentiation time series (Figure 3C). GFP+ and GFP-sorted populations at the starting day 15 time point were easily distinguished based on NKX2-1 transcript expression levels (Figure 3D), validating the efficacy of the NKX2-1^GFP^ reporter. Furthermore, the outgrowth of the GFP+ sorted population could be visualized on SPRING plots as adjacent populations ordered sequentially in time. Apparent bifurcations appeared as multiple branchpoints in transcriptomic trajectories after day 17 (Figure 3C) possibly implying branching cell fates over time, with distinct branches to lung (NKX2-1 positive) and non-lung fates (NKX2-1 negative) such as BAMBI and LGR5, spread over multiple time points (Figure 3D). Trajectories where NKX2-1 expression was maintained after day 17, exhibited subsequent surfactant-encoding and lamellar body-encoding gene expression, beginning on day 21, consistent with a time-dependent alveolar epithelial differentiation program (Figure 3 E-H). We used the mature AEC2 marker profiles identified in primary cells (Figure 1) and we found cells with expression for these markers in late branching parts of the plot representing day 29-31 time points (Figure 3E and H). Eight out of the top 10 most upregulated transcripts in this branch (Figure 3E) were known AEC2 genes that were also present in our primary adult AEC2 differentiation and maturation sets (Figure 1C, D) including SFTPC, CLDN18, CEBPδ, NAPSA, and PGC. Taken together these results are consistent with a fate trajectory followed by a subset of iPSC-derived NKX2-1+ lung progenitors, only some of which reach mature AEC2-like states over a 2-week period.

### A continuous branching network model learns, predicts and maps cell fate paths

Since the SPRING analysis provides a low dimensional dynamic representation implying branching trajectories, we next sought to fully reconstruct these putative branching points to study their regulation and to characterize the set of TFs and signalling pathways associated with their potentially bifurcating fates. For this, we extended our previously developed computational method based on Hidden Markov models (Ding et al., 2018; Rashid et al., 2017) so that it can continuously assign cells along trajectories while still being able to infer regulators controlling branching events. The new method relies on Continuous State Hidden Markov Models (CSHMM). CSHMMs allows us to combine the continuous representation offered by current dimensionality reduction methods with the ability to handle noise, dropouts and identify regulators based on its probabilistic assumptions (Lin and Bar-Joseph, 2019). Unlike standard HMMs which are defined using a discrete set of states, continuous state HMMs can have infinitely many states allowing for continuous assignment of cells along developmental trajectories. A schematic of the learning procedure is depicted in Figure 4A (see Methods for details) resulting in the paths (P0-10) and nodes (N0-11) of a differentiation tree representation, which is not evident when using conventional methods such as tSNE plots (Figure S2).

**Figure 4.**
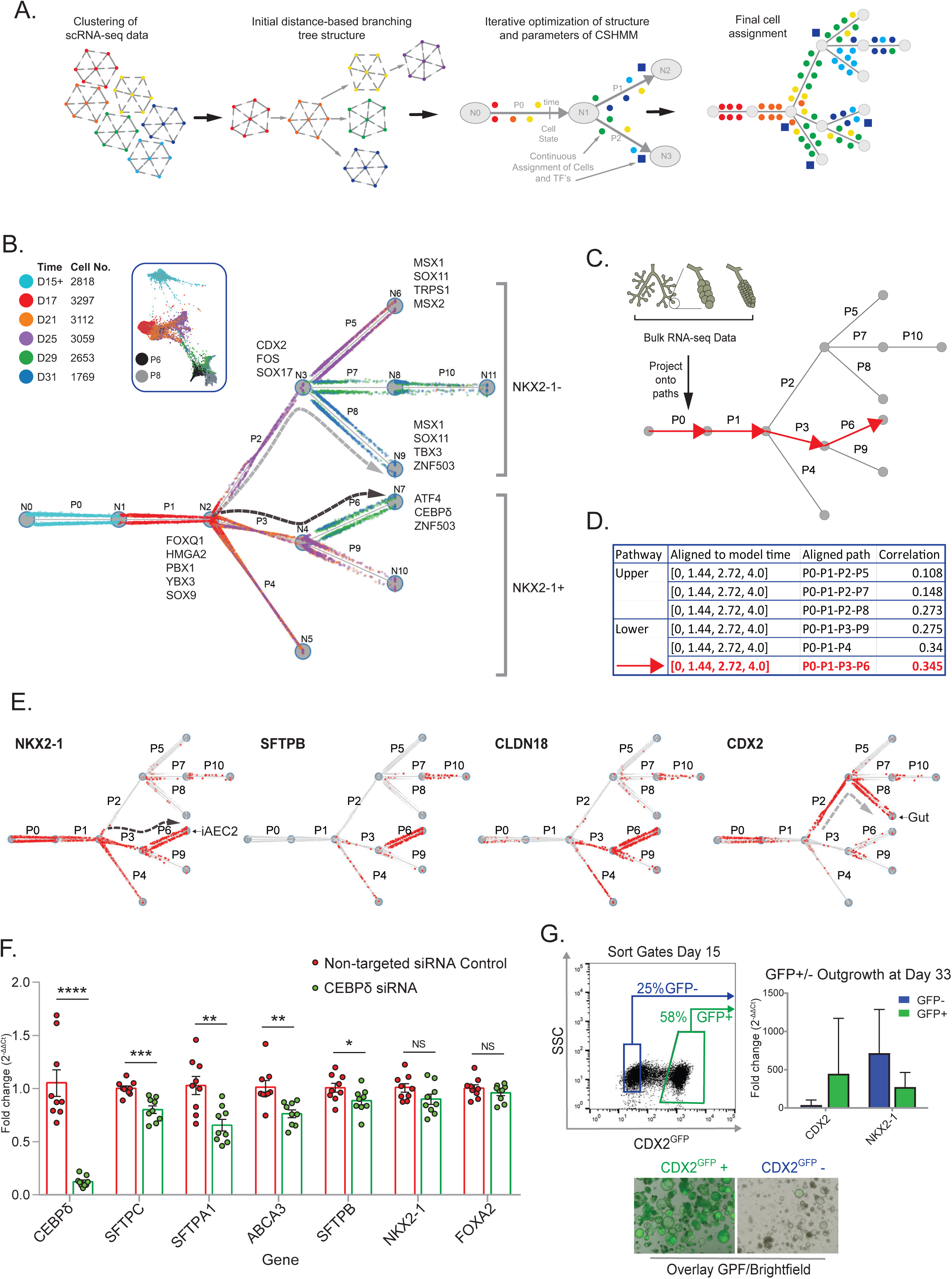
Fate trajectories predicted based on a Continuous State Hidden Markov Model. **(A)** Schematic summarizing the CSHMM method, which starts by clustering cells in each of the time points profiled based on their expression values. Next, clusters are ordered based on distance to the root (first time point cluster). Clusters at each level are linked to the closest cluster at a level above them to generate an initial branching tree. Cells from each cluster are initially assigned to random locations on the branches leading to them. Using these assignments an initial CSHMM model is learned. The model is further refined using an Expectation Maximization (EM) algorithm in which iterations between parameter, model learning, and cell assignment are carried out until convergence. **(B)** The resulting CSHMM model for lung directed differentiation based on scRNA-seq time series data. Each dot represents a cell, color denotes the time point in which the cell was sampled. Nodes are denoted by N0, N1 etc. while branches (paths) are denoted by P0, P1 etc. (note that several branches can share a node). As can be seen, this model predicts that cells remain homogenous in terms of fate commitments until a point between D17 and D21. They then branch to two major paths, an “upper path” containing cells with non-lung endoderm and gut markers, and lower paths (especially P6) that are associated with cells expressing lung markers. Names next to paths are the transcription factors (TFs) that are differentially expressed for these paths. For example, for P6 ATF4 and CEBPδ are significantly differentially expressed TFs whereas for P2 (upper path) CDX2 is significant. (C, D) Alignment of bulk RNA-Seq data from 4 in vivo time points (from Figure 1) to the CSHMM model. The correlation of expression values between the bulk time series data and all possible set of paths in the model was computed. The table lists the correlations for a number of such paths and shows that the best match is for the path that goes from P0 to P6 which is also the path containing the cells with lung epithelial markers. (E) Expression of specific markers in cells assigned by the model to different branches. For example, SFTPB expressing cells are mainly assigned to P6 whereas NKX2-1 cells are assigned to all paths leading to P6 (including to earlier time points). (F) Relative expression levels of each indicated AEC2 or endodermal transcript (RT-qPCR) in iAEC2s after knockdown of CEBPδ by siRNA, validating a role predicted by CSHMM for this TF in regulating the AEC2 program, but not the endodermal (FOXA2) or lung progenitor (NKX2-1) programs. (G) The iPSC line, BU1 with a CDX2^GFP^ reporter was differentiated in triplicate to Day 15, sorted on GFP+ and GFP-cells (representative sort gates shown in first panel) and each outgrowth was cultured in 3D for a further 18 days in the conditions as previously described for distal lung differentiation. At Day 33, cells were harvested, and RT-qPCR was performed on each outgrowth. Relative expression levels of CDX2 and NKX2-1 transcripts are shown in the second panel for each outgrowth showing enrichment for lung and non-lung lineages based on CDX2 and NKX2-1 gene expression. Representative images of microscopy for brightfield and fluorescent overlay of CDX2^GFP^+ and CDX2^GFP^-cells.

To test if the model indeed captures paths corresponding to human lung development, we first compared the reconstructed CSHMM map to our human in vivo expression data by projecting global gene expression from our 4 developmentally relevant time points (Primordial lung progenitor, Early human fetal lung, Late human fetal lung and Adult AEC2, Figure 1) onto the CSHMM map (Figures 4A-C). The best correlation between the model and the in vivo expression data was achieved for the P0-P1-P3-P6 path which leads to AEC2-like cells (Figure 4C lower path). More generally, correlations with lower paths which lead to AEC2-like cells were 20% higher than those with the upper paths indicating that for multiple branches in vitro, PSC-derived differentiation data are in good agreement with in vivo data (Figure 4D). To allow seamless comparisons between CSHMM and SPRING plots we developed an interactive web tool (cosimo.junding.me). Using this tool we found that the P0-P1-P3-P6 CSHMM path matched the implied SPRING trajectory to the mature AEC2 cluster.

Next, we sought to use our reconstructed model to determine when cell fates begin to diverge (Figure 4B). The reconstructed CSHMM model depicted a split in cell fate after day 17, with 6202 cells assigned to the non-lung endoderm (upper) paths and 5980 cells assigned to lung (lower) paths. The upper path, P1-P2-P8 (Figure 4B), was enriched for the expression of intestinal cell marker (CDX2) while the lower path, P1-P3-P6, was enriched for the expression of lung epithelial markers, such as NKX2-1, CLDN18 and SFTPB (Figure 4B, D, and E). Cells in the P10 path were those not assigned to either of these two fates and top differential genes in these cells were mostly mitochondrial genes (see Table S2 for the complete DE gene list for each path). The model identified several transcription factors (TF)s putatively regulating each of the predicted paths at branching points (Figure 4B). TFs assigned to the first major fate split are known regulators for lung epithelial fate including the distal lung developmental regulator SOX9 (Perl et al., 2005; Rockich et al., 2013) and HMGA2, a TF highly expressed in human distal lung bud tip cells (Nikolic et al., 2017) and lung epithelial carcinomas (Snyder et al., 2013). For the lower lung epithelial path to iAEC2 fate (P3-P6, black arrow) the model identified ATF4, CEBPδ and ZNF503 as top TFs most associated with iAEC2 fate, findings in keeping with recent analyses of new-born lungs where CEBPδ and ATF4 are TFs highly expressed in the alveolar epithelium in vivo at post-natal day 1 (Guo et al., 2019). We validated performed siRNA-based knockdown of CEBPδ in iPSC-derived iAEC2s and found this resulted in reduced SFTPC, SFTPB, ABCA3, and SFTPA1 gene expression levels without altering endodermal (FOXA2) or lung TF (NKX2-1) levels (Figure 4F) consistent with a role for CEBPδ in maintenance of the AEC2-specific program. The branching model also predicted that CDX2, FOS, and SOX17 are top TFs associated with the non-lung endoderm or intestinal fate paths (P2-P8, grey arrow), and we validated this finding in independent experiments using another iPSC line (BU1 carrying a CDX2^GFP^ knock-in reporter; Mithal et al. 2018, Manuscript in submission) observing that sorted CDX2^GFP+^ cells at day 15 of our lung differentiation protocol were markedly enriched in intestinal competence, in contrast to CDX2^GFP^ negative cells which were enriched for lung competence and depleted for intestinal competence (Figure 4G).

### CSHMM predicts the precise timing of Wnt modulation that maintains lung cell fate

In addition to TFs, the branching identified by the CSHMM model assigned cells in which the Wnt and BMP signaling pathways were upregulated in the progeny of sorted NKX2-1^GFP+^ cells as they diverged to the non-lung paths (Figure 5A). Specifically, 4 of the 5 top differentially expressed genes in the non-lung endodermal (upper) path were related to Wnt signaling: WIF (Ng et al., 2014), HIPK2 (Tan et al., 2014), NEAT1 (Zarkou et al., 2018), and THBS1 (Han et al., 2014) (Table S3). Furthermore, Wnt target genes LEF1, NKD1, and AXIN2 (McCauley et al., 2017) were all upregulated in cells following non-lung paths, compared to those maintaining lung paths. We and others have observed that downregulation of Wnt signalling targets has stage-dependent effects in lung development in vivo (Frank et al., 2016; Mucenski et al., 2003; Shu et al., 2005) and in vitro (Jacob et al., 2017; McCauley et al., 2017), inducing proximal airway patterning when downregulated at the NKX2-1+ primordial progenitor stage (PSC differentiation day 15) whereas downregulation in distal lung epithelium *in vivo* (Frank et al., 2016) or in iAEC2s (Jacob et al., 2017) at later stages is associated with distal lung maturation, as validated in our human primary cell RNA-seq (Figure 1). However, the optimal timing of downregulation of Wnt, for example by withdrawal of the GSK3 inhibitor, CHIR, in our system has not been established. Therefore, we used the reconstructed branching model to predict the optimal time point for Wnt withdrawal in order to maximize the set of cells maintaining lung fate in our protocol. To find that point we selected a set of canonical Wnt signalling target genes and plotted their expression in the lung and non-lung trajectories. As can be seen in Figure 5B, these genes start to diverge at the mid-point of P1. To assign an actual time to that point we looked at cells assigned by CSHMM before and after that midpoint and computed the average time in which these cells were profiled. Using this, we determined that day 17.5 (Red arrow in Figure 5B) is the time of split between the two branches. To test this prediction, we repeated our directed differentiations while withdrawing CHIR from our media for a period of 4 days (Figure 5C), starting at five different time points over the 2 week period of differentiation of sorted NKX2-1^GFP^+ lung progenitors towards the desired iAEC2 target (days 15-29). To maintain proliferation of resulting cells, CHIR was added back after 4 days, allowing each parallel condition to be harvested at the identical total differentiation time while keeping the length of CHIR withdrawal (4 days) constant for each condition. As predicted by the model, withdrawal of CHIR beginning on day 17 resulted in the highest rates of retention of distal lung epithelial fate as quantified by flow cytometry measurement of NKX2-1^GFP^ and SFTPC^tdTomato^ reporter expression on day 29 (Figure 5D and E), Overall, these experiments demonstrate that CSHMM can not only identify the relevant signalling pathways which determine cell fate in our system, but can also predict with precision the timing of pathway modulation to increase differentiation efficiency to the target cell.

**Figure 5.**
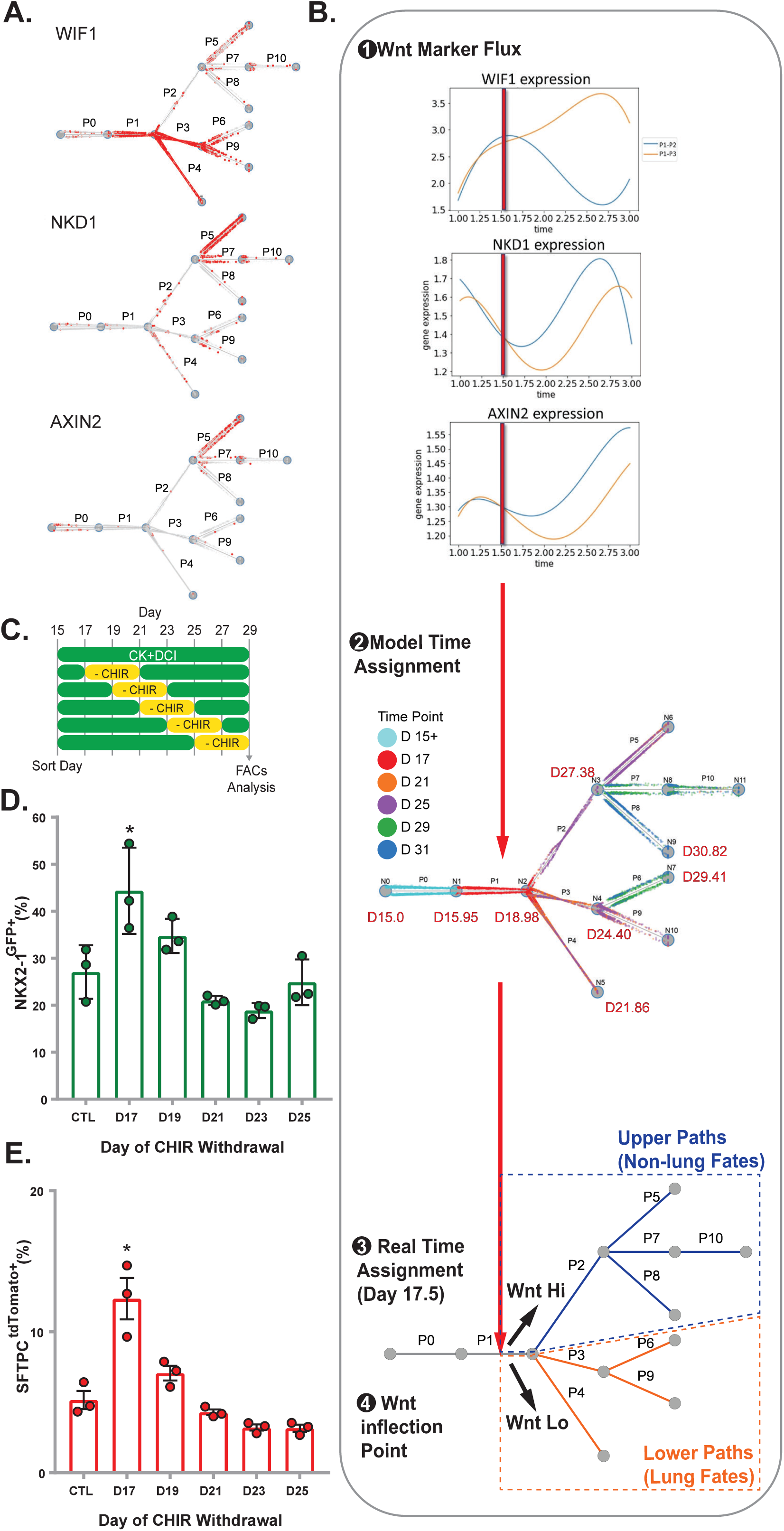
CSHMM predicts the precise timing of Wnt modulation as a determinant of cell fate. (A) Expression of key Wnt target genes enriched in upper paths, whereas Wnt inhibitory factor, WIF1, is enriched in lower paths. (B) To determine the exact time of Wnt pathway activation the continuous expression of these markers is reconstructed using splines to plot the reconstructed expression profiles for the three markers for cells assigned to the top paths (blue curve) vs. bottom paths (orange curve). For all three there is a split in expression values at the halfway point between nodes N1 and N2 (middle of P1). To determine the real time denoted by this point a time is assigned for each node in the CSHMM tree by averaging the profiled times for cells assigned right before and right after this node. Since the two nodes that define P1 are assigned times D15.95 and D18.98 respectively, the middle point between them is D17.5, the predicted split time. (C) Schematic summarizing experimental plan for testing effect of time-dependent downregulation of canonical Wnt signalling by CHIR withdrawal. (D) Retention of distal lung epithelial fate on day 29 of the experiment in C, measured by the frequency of cells expressing the NKX2-1^GFP^ and SFTPC^tdTomato^ reporters quantified by FACS. *=ANOVA p<0.05.

### Lineage tracing using DNA barcoding reveals clonal heterogeneity and fate plasticity

The CSHMM computationally predicts multipotency at least until day 17.5, with some cells branching to lung and others to non-lung after this time. To functionally test this prediction, we employed lentiviral barcoding to clonally trace the progeny of individual cells in the protocol followed by scRNA-seq profiling to assign them to paths in the model. On day 15 of differentiation PSC-derived NKX2-1 progenitors were sorted to purity and on day 17 a single cell suspension of these progenitors was infected with our lentiviral barcoding library for “Lineage and RNA Recovery” (LARRY) (Weinreb et al., 2019) which encodes for enhanced green fluorescent protein (eGFP) mRNA together with a 3’UTR carrying a unique inheritable barcode for each cell (Figures 6A). This library has a complexity of 10^6^ barcodes, sufficient to label 10^4^ cells with <0.5% barcode overlap between clones (Weinreb et al., 2019). We first optimized this system to achieve a transduction efficiency in iPSC-derived lung progenitors of ∼30% using a viral multiplicity of infection (MOI)=10. Following lentiviral infection of 32,500 progenitors on day 17, both the infected cells and parallel uninfected (MOI=0) control progenitors were cultured for an additional 10 days in our distal lung media prior to capture for scRNA-seq (Figure 6A). Analysis of the single cell transcriptomes of 6,147 cells from the MOI=10 condition and 1644 cells from the MOI=0 control revealed four cell states or clusters (Figure 6B). Comparing infected (MOI=10) to uninfected (MOI=0) samples by tSNE plots revealed overlay of all 4 clusters, indicating that lentiviral infection and tagging did not detectably perturb differentiation (Figure S3). We annotated the 4 cell clusters, based on the expression of identity marker genes as distal lung epithelium (hereafter “lung”) and “non-lung endoderm”; and 2 minor clusters: pulmonary neuroendocrine cells (“PNEC”) and “gut”, (Figure 6B and C, Table S4 for full list).

**Figure 6.**
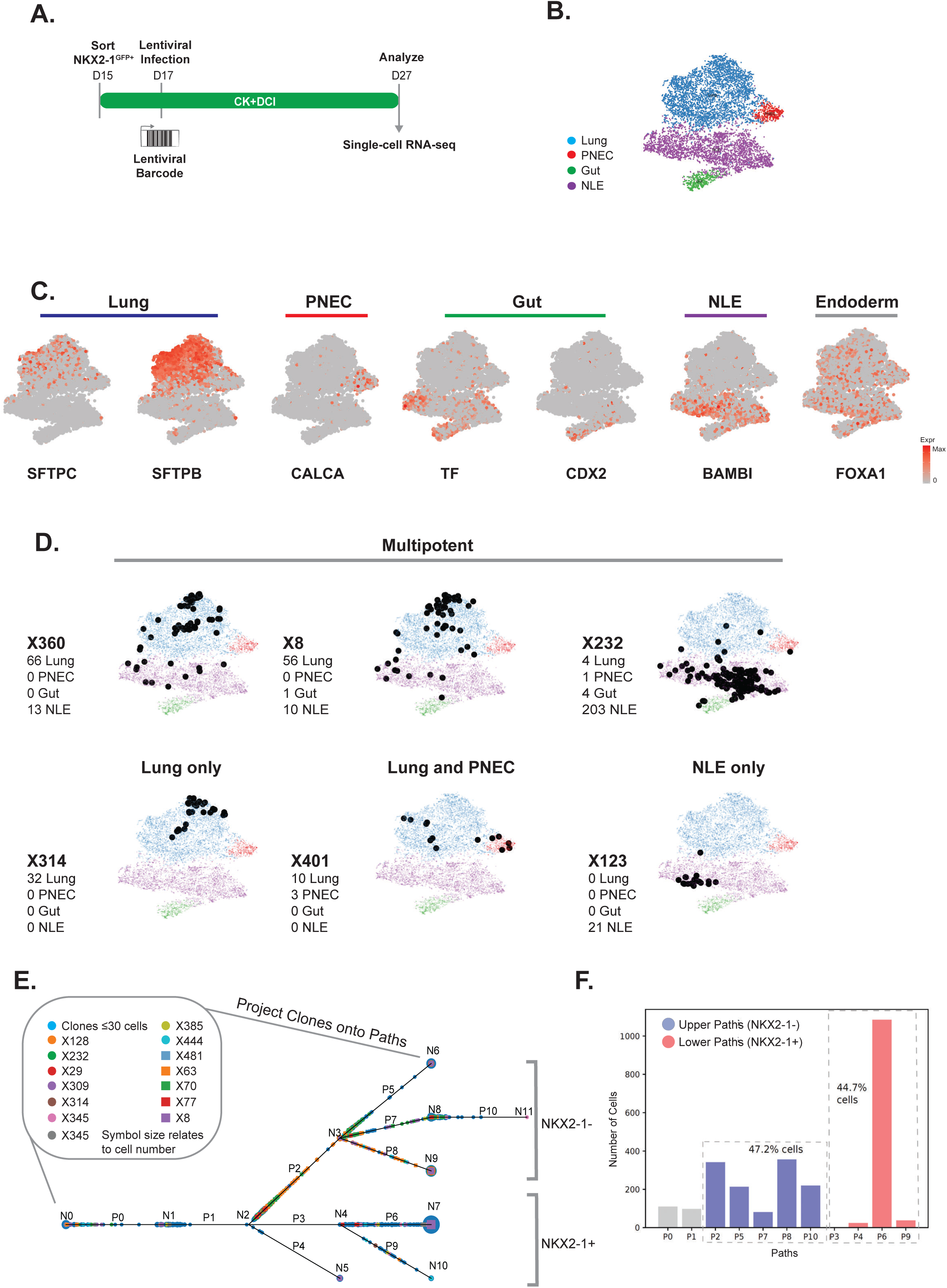
Lineage tracing using lentiviral barcoding reveals clonal heterogeneity. (A) Schematic of experiment showing infection of NKX2-1 + outgrowth at day 17 with lentivirus to tag progenitors with unique integrated DNA barcodes. After infection cells were replated in 3D Matrigel and grown for a further 10 days in CK+DCI media at which point a single cell suspension was generated and cells were encapsulated for scRNA-seq using inDrops. Inherited lentiviral barcodes were matched with transcriptomic profiles for each cell to track clones from day 17 to day 27. (B) tSNE plot of all cells harvested at day 27 with Louvain clustering (resolution 0.25) which has been annotated based on marker genes for distal lung alveolar epithelium ‘Lung’, pulmonary neuroendocrine cells (PNEC), Gut and non-lung endoderm (NLE). (C) Normalized gene expression overlayed on tSNE plots for selected markers of ‘Lung’ (SFTPB and SFTPC), ‘PNEC’ (CALCA), ‘Gut’ (TF and CDX2), ‘NLE’ (BAMBI) and retention of endoderm lineages (FOXA1). (D) tSNE plots with Louvain clustering for annotated cell lineage and overlayed clones X360, X8 and X232 which are found contributing to multiple cell lineages labelled ‘Multipotent’. Also overlayed are examples of clones found in ‘Lung’ lineage only (X314), ‘Lung’ and ‘PNEC’ (X401) and NLE only (X123). (E) Lentivirally barcoded cells projected onto the CSHMM based on the expression of 86 selected genes. Cells are colored based on individual lentibarcodes indicating clones arising from distinctly tagged individual ancestors. Note: several large clusters are assigned to both top and bottom paths, validating the bifurcating trajectories predicted by the CSHMM and indicating that cell fate is not fully determined by D17. (F) Percentage of lentibarcoded cells assigned to top and bottom paths. Similar proportions of cells are assigned to the paths as were seen in the original dataset (without lentiviral infection) indicating that the insertion of the virus did not appreciably impact or bias the differentiation of cells.

We identified all lentivirally transduced clones within each cell cluster by associating lentiviral barcodes to cell transcriptomes (Figure 6 and Table S5). We identified 487 unique clones with 45 clones containing more than 10 cells per clone (Figure S3). The majority of these 45 clones contained cells which were found in more than one cell state (23/45 contributing to both lung (lung or PNEC) and non-lung (NLE or gut) clusters; Figure 6D and Figure S3). For example, the largest clone X232 (212 cells) was found to contribute progeny to all 4 cell clusters implying that at least a subset of day 17 parental progenitors was likely following the bifurcating tree predicted by CSHMM.

We next directly overlaid barcoded cells on the CSHMM fate maps (Figure 6E). Given experimental differences in profiling cells used to construct the CSHMM model and barcoded cells we projected barcoded cells using a subset of 82 genes, including the top 68 genes differentially expressed between the upper and lower paths (P2 and P3) and 14 distal lung markers from our primary cell datasets (Table S6). Randomization analysis showed that projections based on the set of 82 genes led to significant correlations between the initial and barcode scRNA-seq levels (based on both Ranskum and t-tests; Table S7). Using these genes for the projections we found that only a few cells were assigned to the earlier paths (P0 and P1) whereas over 90% of cells were correctly assigned to the later paths (after P0 and P1). As for the two major branches (Upper, non-lung and Lower, lung) we found that 13/14 (92.9%) of the largest clones (>30 cells) were assigned to both paths and 108/272 of all clones (39.7%) with > 1 cell were assigned to both upper and lower paths (Table S6). Projecting the largest clones (≥ 30 cells) on cell fate paths predicted by CSHMM (Figure 6E and F) suggested that no predominant clone contributed uniquely to each cell fate path confirming that cells may still switch cell fate after day 17.

While the two complementary analysis methods we used, tSNE cluster assignments (unsupervised) and CSHMM projections (supervised) led to the same conclusions about multi-potency, we further examined the agreement of each method for each of the largest clones (>30 cells). We found overall good agreement between the way the largest clones were assigned by the two methods (Table S8). Taken together, these results indicate that DNA barcoding agrees with the CSHMM prediction that cell fate is not completely decided before day 17 and therefore PSC-derived progenitors are still “plastic” or multipotent at this developmental stage.

### Time dependent maturation results in stabilization of lung epithelial cell fates allowing indefinite expansion of iAEC2s in culture

Our detailed model covered the period between D15 and D31 and identified several branching events and their regulation with a specific path leading to the desired iAEC2 phenotype. We hypothesized that the frequency of reversion to non-lung endoderm, observed in the model, might decline over time as cells mature, allowing the propagation in culture of iPSC-derived lung cells with more stable phenotypes (Figure 7A). To test for this possibility, we evaluated RUES2 ESCs, BU3 NGST iPSCs, as well as two additional iPSC lines (SPC2 and ABCA35) each targeted with our SFTPC^tdTomato^ reporter to allow real time monitoring of distal lung fate. Differentiating each line via purified lung progenitors (sorted on day 15) again resulted in a day 30 cell population that contained mixed lung (NKX2-1+) and non-lung endoderm (NKX2-1-). However, resorting this population based on SFTPC^tdTomato^ expression after this period (day 51) (Figure 7B, C), resulted in the outgrowth of epithelial spheres that maintained SFTPC^tdTomato^ expression indefinitely, as we have previously published for RUES2 and BU3 cell lines (Jacob et al., 2017). We validated this same pattern for SPC2 and ABCA35 iPSCs, finding each line maintained SFTPC^tdTomato^ expression in >95% of cells followed as serially passaged cell cultures for at least 64 days and 102 days after sorting SFTPC^tdTomato^ + cells on days 51 or 34 of differentiation, respectively (3-4 passages post sorting; Figure 7C-E and Figure S5; total differentiation time 115 and 136 days respectively). scRNA-seq of the outgrowths from each line were prepared without further purification, and tSNE visualizations validated retention of distal lung phenotype in almost all cells without reversion to non-lung endoderm (Figure 7F-H and Figure S5). For example, <0.1% of SPC2 derived cells at this time point expressed the gut marker CDX2, or hepatic markers TF, AFP, or ALB, and only 2 out of 1390 cells expressed gastric marker TFF1 (Figure 7H). All cell clusters expressed high levels of NKX2-1 and AEC2 markers (Figure 7G) with similar findings in ABCA35 iPSC-derived cells (Figure S5). Based on this stability in iAEC2 phenotype, using this approach we could generate 10^20 iAEC2s per input sorted tdTomato+ cell over a 225 day period without further cell sorting. Taken together these results suggested that as in other in vivo developmental systems, there is time dependent loss of plasticity and stabilization or restriction of cell lineage in our model system.

**Figure 7.**
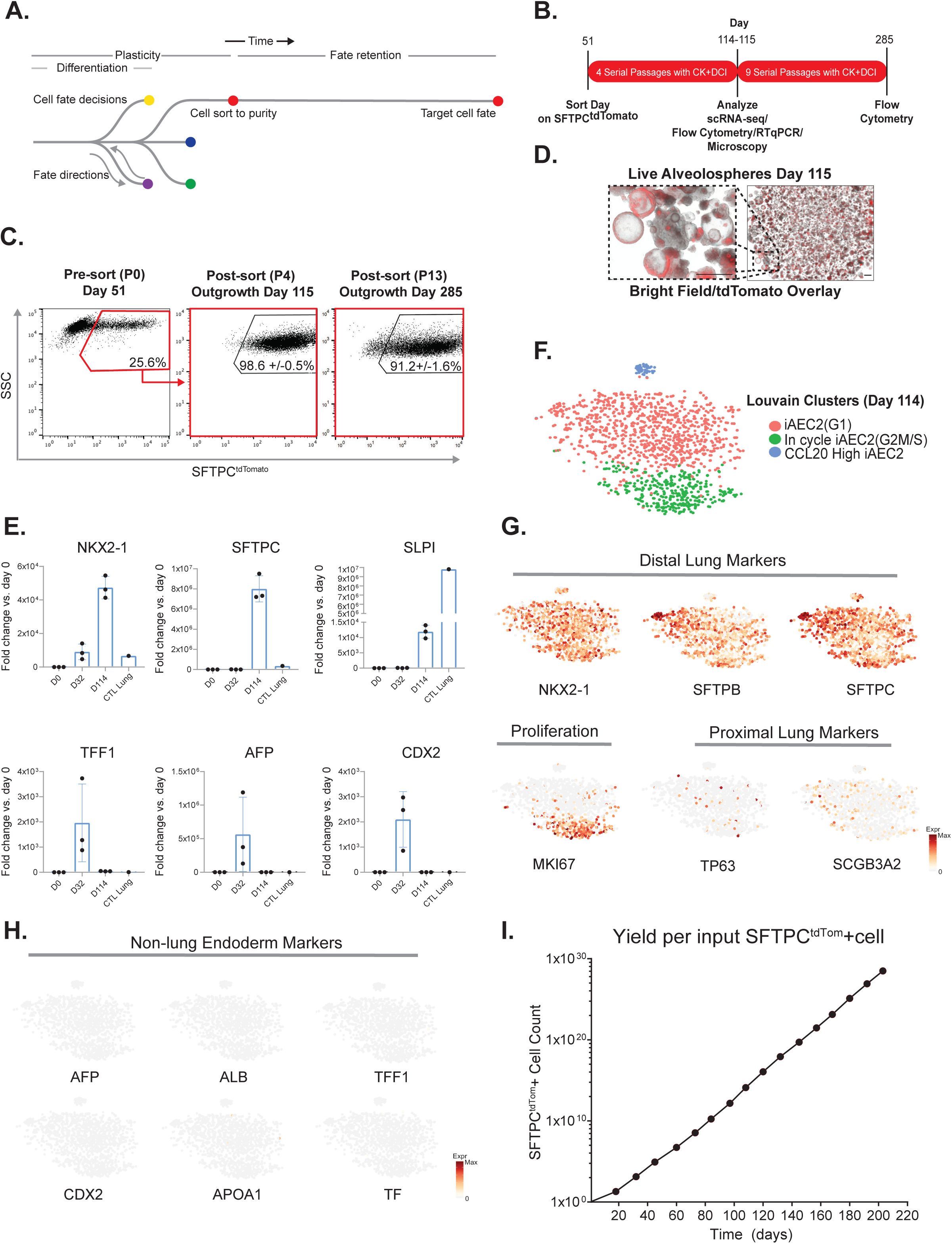
Time Dependent Maturation Leads to Stabilized Lung Fate Retention of iAEC2. (A) Overview schematic of differentiation, maturation and fate retention after mature cells are sorted to purity and further cultured for extended periods of time without loss of lung cell fate. (B) Schematic of experiment in which cells sorted at day 51 for SFTPC^tdTomato^ were replated in 3D conditions in CK+DCI media. They were subsequently passaged as single cells 4 times on the days indicated, without further cell sorting. At day 114, cells were isolated for RT pPCR and at day 115 all live cells were encapsulated for scRNA-seq using the 10X Chromium System. Remaining cells were cultured for a further 170 days to day 285 when flow cytometry was performed. (C) Flow cytometry dot plots of cells before sorting for SFTPC^tdTomato^ and after replating tdTomato+ cells for outgrowth as alveolospheres. Repeated flow cytometry was performed after 4 passages (P4) on day 115 without additional sorting, immediately before scRNA-seq. Almost all cells have retained expression of the SFTPC lung reporter (mean +/-SD is indicated; n=3 biological replicates). Remaining outgrowth cells were cultured without further sorting with repeated flow cytometry performed after 9 further passages (P13) at day 285. (D) Representative images of live SPC2 alveolospheres (bright-field/tdTomato overlay; day 115; at 4X and 20X) illustrating retention of lung fate, indicated by continued expression of SFTPC^tdTomato^. Scale bar, 500 µm. (E) Relative expression levels of each indicated AEC2 or endodermal marker transcript (RT-qPCR) in iAEC2s at day 0 (D0), at day 32 (D32) before sorting for a pure population of SFTPC^tdTomato^ and at day 114 (D114; four passages after tdTomato+ sorting and extended culture.) Control samples are an adult human distal lung explant (CTL Adult Lung). (F) tSNE with Louvain Clusters annotated using marker genes for PSC-derived AEC2 ‘iAEC2’, markers of cells in the G2/MS cell cycle stages ‘In cycle iAEC2(G2/MS)’ and iAEC2 with high CCL20 ‘CCL20 High iAEC2’. (G) Normalized gene expression overlayed on tSNE plots for selected markers for ‘Distal Lung’ (NKX2-1, SFTPB, and SFTPC), ‘Proliferation’ (MKI67), ‘Proximal Lung’ (TP63 and SCGB3A2), or (H) ‘Non-Lung Endoderm’ (AFP, ALB, TFF1, CDX2, APOA1 and TF). (I) Line graph of yield per input SFTPC^tdTomato^+ cell. In a separate differentiation experiment, SPC2 cells were sorted for SFTPC^tdTomato^ + cells at day 45 and the positive outgrowth was cultured with 15 passages for a further 203 days to day 248. At each passage and at day 248 flow cytometry was performed for SFTPC^tdTomato^ and positive cells per well were counted (n=3). The line plot represents the cumulative count of cells per input cell shown on the y-axis with time on the x-axis, and demonstrates that the system can generate 10^20 iAEC2s per input sorted tdTomato+ cell over a 225 day period without further cell sorting.

## Discussion

To improve our understanding of PSC differentiation protocols we developed a new framework that combines experimental design, computational modelling, lentiviral barcoding, and scRNA-seq profiling. We first established human fetal and adult primary cell datasets to serve as benchmarks against which in vitro engineered cells can be compared, and we used our method to map the dynamic changes that occur as lung progenitors differentiate towards mature AEC2s. The reconstructed continuous branching models generated by CSHMM provide details about developmental paths, branching, and regulating TFs. Each point in the model can be mapped back to real time enabling the prediction of time specific interventions. We validated several predictions of the model including several predicted TFs for specific paths, the multipotency it implies, and the timing of a predicted intervention that leads to better retention of the desired cell fate.

As is evident by tracing descendants of lentivirally barcoded parents, clonal plasticity is observed in our PSC-derived system leading to lung and non-lung endodermal cell fates, a finding which parallels in vivo observations where developing or adult lung epithelia tend to revert to non-lung endodermal fates in abnormal or diseased settings where exuberant Wnt activity is present or where there is loss of Nkx2-1(Herriges et al., 2014; Okubo and Hogan, 2004; Snyder et al., 2013; Tata et al., 2018). For example, Hogan and colleagues found non-lung endoderm, such as intestinal programs, emerged in mouse lung epithelial cells in vivo after lineage-specific conditional hyperactivation of canonical Wnt signalling (Okubo and Hogan, 2004). In addition, in settings where Nkx2-1 expression is lost, multiple lung epithelia revert to non-lung endodermal fates in either fetal or adult lungs through a mechanism that involves loss of repression of Foxa2-driven non-lung fates (Snyder et al., 2013). This emergence of non-lung endodermal descendants from lung epithelial parents is particularly evident in lung adenocarcinoma settings (Snyder et al., 2013; Tata et al., 2018), suggesting that our in vitro model may provide insights for understanding and preventing the fate changes that occur during lung cancer pathogenesis.

We validated the parent-progeny relationships predicted by our model by using a combination of scRNA-seq and DNA barcoding (Weinreb et al., 2019). Genetic tagging of individual cells allowed tracing of their progeny during directed differentiation. Such an approach can match lineage relationships in both a supervised and unsupervised manner, as has been recently reported (Biddy et al., 2018; Wagner et al., 2018). Finally, we also found that cells that are assigned to the AEC2 path in our model appear to stabilize their phenotypes, consistent with in vivo patterns of time dependent restrictions of developing fates including endodermal lineages (Grapin-Botton, 2005). This leads to an expandable pool of lung progenitors that maintain stable AEC2-like fate even after extensive proliferation in vitro.

Several limitations to our methods are important to highlight. While we sampled cells at the earliest possible time after sorting for progenitors, earlier acquisition of endodermal samples before sorting on the lung progenitor marker NKX2-1 or the use of epigenetic profiling could identify pre-patterning of fates that might have been missed in our transcriptomic profiles. Second, our method was designed to detect multipotency and fate bifurcations rather than to quantify any lineage bias that may be present at each developmental stage. Others have published lentiviral barcoding over time that might be employed to more precisely quantify lineage bias at each stage in our protocol (Biddy et al., 2018; Weinreb et al., 2019). It should be pointed out that prior time series profiles have revealed fate convergence from distinct origins is detectable in the development of alternate germ layer derivatives (e.g. neural crest cells; (Wagner et al., 2018)), however, we found only divergence rather than convergence to be present in our lung developmental trajectories, suggesting convergence may not contribute to the emergence of AEC2s. These results are consistent with prior observations that all distal lung epithelial descendants arise via the gateway of an endodermal NKX2-1+ progenitor, rather than from alternate origins (Hawkins et al., 2017; Jacob et al., 2017; Longmire et al., 2012).

Despite these limitations, the framework we developed which combines predictive computational approaches with cell fate tracing is generalizable. It can be used to further understand and model several other directed differentiation strategies and disease pathogenesis, potentially leading to future cell therapies.

## Acknowledgments

The authors wish to thank Caleb Weinreb and Allon Klein of Harvard Medical School for assistance with their lentiviral barcoding library, the LARRY pipeline, access to the HMS inDrops core facility and application of SPRING software. We thank Yuriy Alekseyev of the Boston University School of Medicine (BUSM) Single Cell Sequencing Core and Brian R. Tilton of the BUSM Flow Cytometry Core; both supported by NIH grant 1UL1TR001430. For facilities management, we are indebted to Greg Miller, CReM Laboratory Manager, and Marianne James, CReM iPSC Core Manager, supported by grants R24HL123828 and U01TR001810. We are grateful to Michael Morley at the University of Pennsylvania for ongoing access to his bioinformatics portal for analyses of bulk RNA-seq datasets. The current work was supported by grants from the Alpha-1 Foundation (KH), TL1TR001410 and F31HL134274 (AJ), F30HL142169 (YS), the I.M. Rosenzweig Junior Investigator Award from the Pulmonary Fibrosis Foundation (KDA), R01GM122096 and R01HL128172 (ZBJ and DNK), 0T20D026682 (ZBJ), and R01HL095993, R01HL122442, U01HL134745, and U01HL134766 (DNK).

## Author Contributions

K.H., D.N.K., Z.B.J and J.D. designed the project, performed experiments and bioinformatic analysis, and wrote the manuscript. K.H., M.J.H., K.D.A, Y.S., R.B.W., A.M., A.J. and M.V. performed differentiation experiments and K.H., M.J.H., K.D.A, Y.S., performed scRNA-seq experiments. Z.B.J and J.D. developed CSHMM. C.V.M. and N.C. analysed scRNA-seq and RNA-seq experiments and performed statistical analysis. S.H.G. provided primary fetal lung samples which were analysed by A.J. by bulk RNA-seq. F.A. and N.K. performed NanoString mRNA quantification. A.R.F and F.C. developed and provided barcoding lentivirus.

## Star Methods

### Key Resources Tables

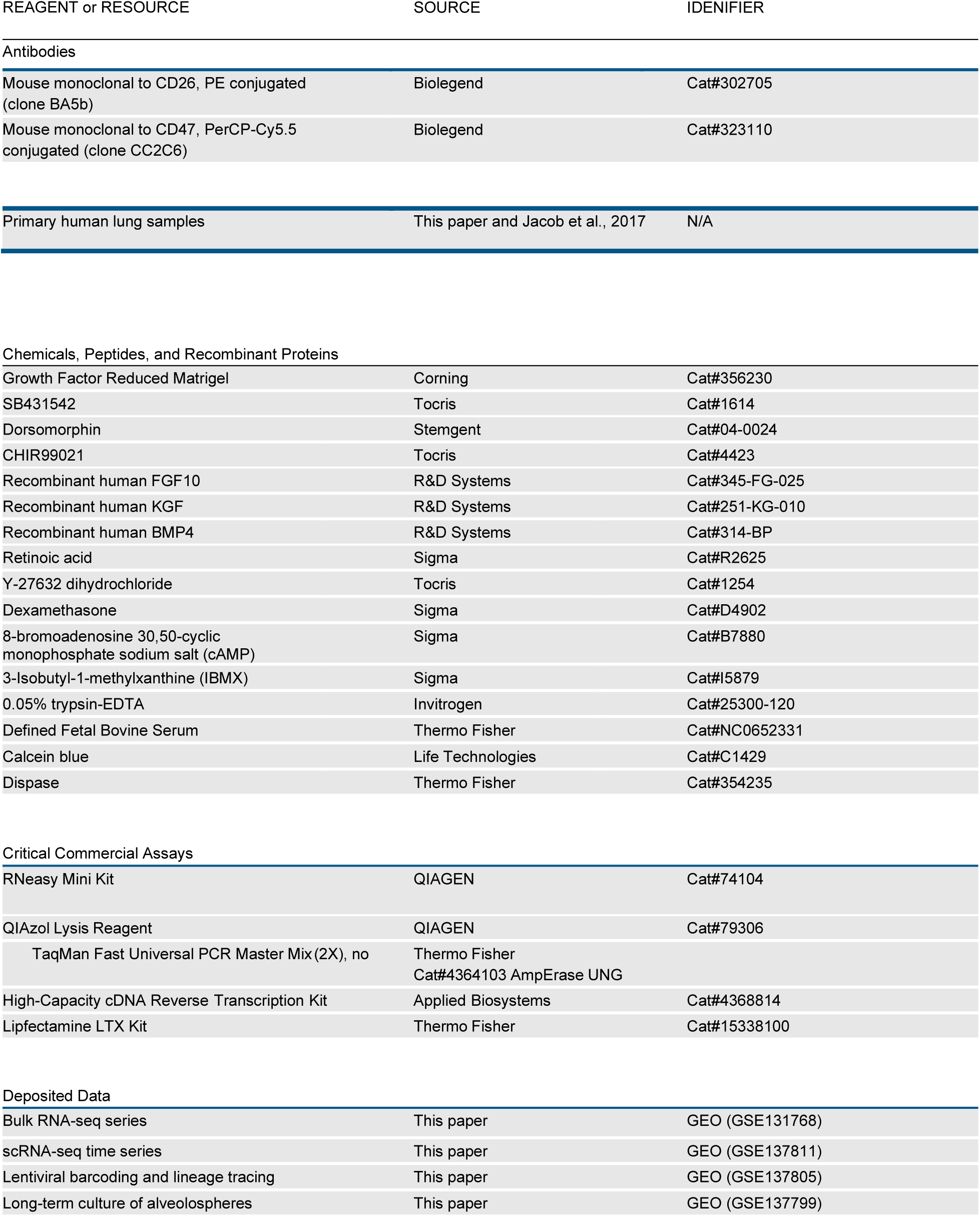
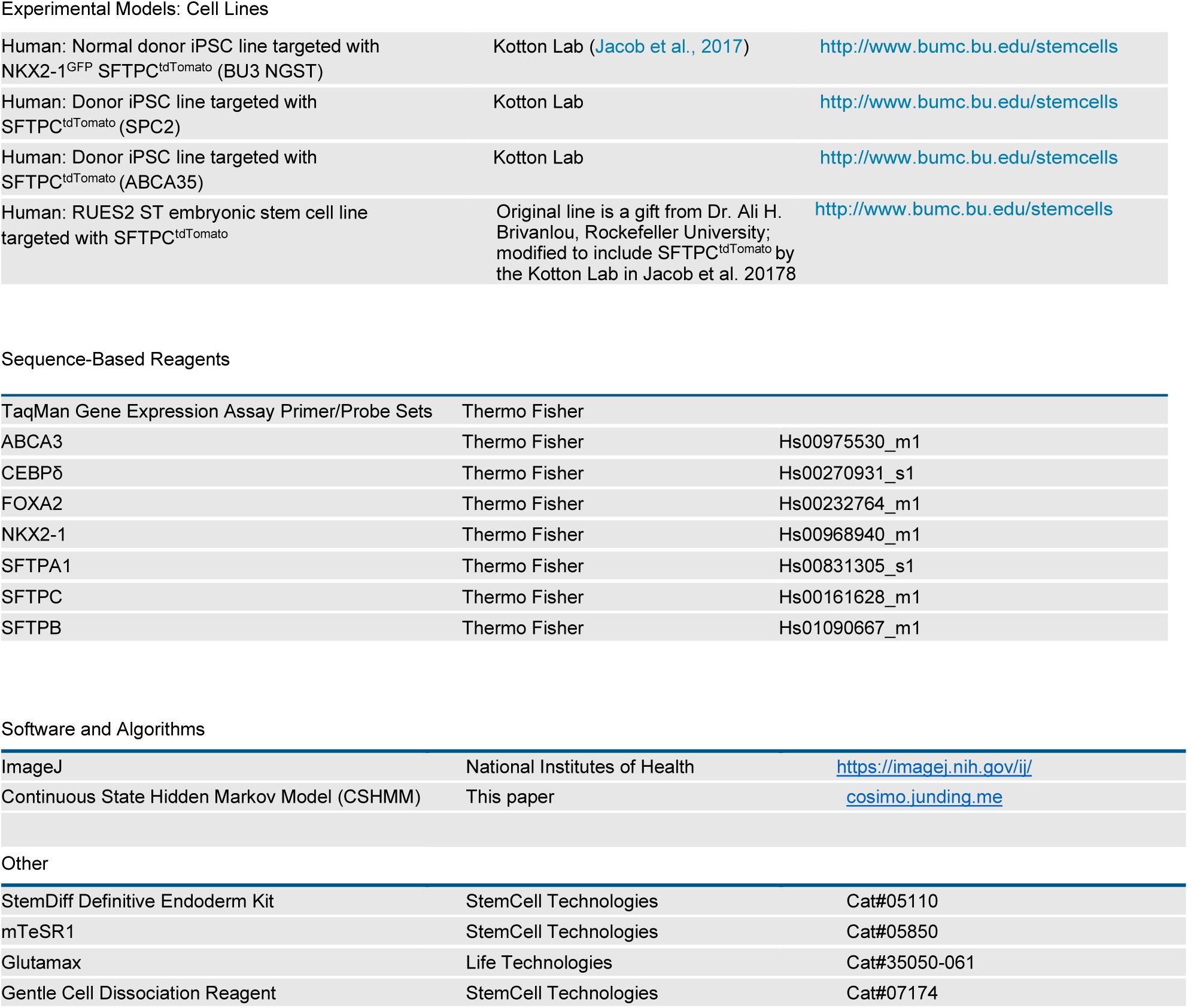

| REAGENT or RESOURCE | SOURCE | IDENTIFIER |
| --- | --- | --- |
| **Antibodies** |  |  |
| Mouse monoclonal to CD26, PE conjugated (clone BA5b) | Biolegend | Cat#302705 |
| Mouse monoclonal to CD47, PerCP-Cy5.5 conjugated (clone CC2C6) | Biolegend | Cat#323110 |
| Primary human lung samples | This paper and Jacob et al., 2017 | N/A |
| **Chemicals, Peptides, and Recombinant Proteins** |  |  |
| Growth Factor Reduced Matrigel | Corning | Cat#356230 |
| SB431542 | Tocris | Cat#1614 |
| Dorsomorphin | Stemgent | Cat#04-0024 |
| CHIR99021 | Tocris | Cat#4423 |
| Recombinant human FGF10 | R&D Systems | Cat#345-FG-025 |
| Recombinant human KGF | R&D Systems | Cat#251-KG-010 |
| Recombinant human BMP4 | R&D Systems | Cat#314-BP |
| Retinoic acid | Sigma | Cat#R2625 |
| Y-27632 dihydrochloride | Tocris | Cat#1254 |
| Dexamethasone | Sigma | Cat#D4902 |
| 8-bromoadenosine 30,50-cyclic monophosphate sodium salt (cAMP) | Sigma | Cat#B7880 |
| 3-Isobutyl-1-methylxanthine (IBMX) | Sigma | Cat#I5879 |
| 0.05% trypsin-EDTA | Invitrogen | Cat#25300-120 |
| Defined Fetal Bovine Serum | Thermo Fisher | Cat#NC0652331 |
| Calcein blue | Life Technologies | Cat#C1429 |
| Dispase | Thermo Fisher | Cat#354235 |
| **Critical Commercial Assays** |  |  |
| RNeasy Mini Kit | QIAGEN | Cat#74104 |
| QIAzol Lysis Reagent | QIAGEN | Cat#79306 |
| TaqMan Fast Universal PCR Master Mix (2X), no | Thermo Fisher | Cat#4364103 AmpErase UNG |
| High-Capacity cDNA Reverse Transcription Kit | Applied Biosystems | Cat#4368814 |
| Lipfectamine LTX Kit | Thermo Fisher | Cat#15338100 |
| **Deposited Data** |  |  |
| Bulk RNA-seq series | This paper | GEO (GSE131768) |
| scRNA-seq time series | This paper | GEO (GSE137811) |
| Lentiviral barcoding and lineage tracing | This paper | GEO (GSE137805) |
| Long-term culture of alveolospheres | This paper | GEO (GSE137799) |
| Experimental Models: Cell Lines |  |  |
| Human: Normal donor iPSC line targeted with NKX2-1<sup>GFP</sup> SFTPC<sup>tdTomato</sup> (BU3 NGST) | Kotton Lab (Jacob et al., 2017) | http://www.bumc.bu.edu/stemcells |
| Human: Donor iPSC line targeted with SFTPC<sup>tdTomato</sup> (SPC2) | Kotton Lab | http://www.bumc.bu.edu/stemcells |
| Human: Donor iPSC line targeted with SFTPC<sup>tdTomato</sup> (ABCA35) | Kotton Lab | http://www.bumc.bu.edu/stemcells |
| Human: RUES2 ST embryonic stem cell line targeted with SFTPC<sup>tdTomato</sup> | Original line is a gift from Dr. Ali H. Brivanlou, Rockefeller University; modified to include SFTPC<sup>tdTomato</sup> by the Kotton Lab in Jacob et al. 2017 | http://www.bumc.bu.edu/stemcells |

Sequence-Based Reagents
| TaqMan Gene Expression Assay Primer/Probe Sets | Thermo Fisher |  |
| --- | --- | --- |
| ABCA3 | Thermo Fisher | Hs00975530_m1 |
| CEBPδ | Thermo Fisher | Hs00270931_s1 |
| FOXA2 | Thermo Fisher | Hs00232764_m1 |
| NKX2-1 | Thermo Fisher | Hs00968940_m1 |
| SFTPA1 | Thermo Fisher | Hs00831305_s1 |
| SFTPC | Thermo Fisher | Hs00161628_m1 |
| SFTPB | Thermo Fisher | Hs01090667_m1 |
| Software and Algorithms |  |  |
| ImageJ | National Institutes of Health | https://imagej.nih.gov/ij/ |
| Continuous State Hidden Markov Model (CSHMM) | This paper | cosimo.junding.me |
| Other |  |  |
| StemDiff Definitive Endoderm Kit | StemCell Technologies | Cat#05110 |
| mTeSR1 | StemCell Technologies | Cat#05850 |
| Glutamax | Life Technologies | Cat#35050-061 |
| Gentle Cell Dissociation Reagent | StemCell Technologies | Cat#07174 |

### Contact#for Reagent and Resource Sharing

Further information and requests for reagents may be directed to, and will be fulfilled by, the Lead Contact, Darrell Kotton.

### Analysis of Primary Human Fetal Lung Time Series by Bulk RNA Sequencing

#### Isolation of primary fetal and adult AECs

Primary fetal lung alveolar epithelial cells and adult AEC2s were isolated for RNA extraction and analysis by bulk RNA-seq as detailed in our prior publication (Jacob et al., 2017) with partial datasets (for only the 21 week and adult cells) previously published in that manuscript and now re-deposited with the Gene Expression Omnibus (GEO) under GSE131768. For the present study 3 additional unpublished fetal alveolar epithelial samples from weeks 16-17.5 of gestation and 1 additional sample from week 20 of gestation was added for analysis. In order to avoid technical batch effects, all 13 samples for bulk RNA sequencing (day 15 PSC-derived primordial progenitors; n=3 biological replicates; week 16-17.5 alveolar epithelium; n=3; week 20-21 alveolar epithelium; n=4, and adult AEC2s; n=3 donor lungs) underwent simultaneous parallel extraction of RNA, library preparation, and RNA sequencing. In brief each sample was isolated as follows: fetal lung tissue (weeks 16-21) was obtained in the Guttentag laboratory under protocols originally reviewed by the Institutional Review Board at the Children’s Hospital of Philadelphia and subsequently reviewed by Vanderbilt University. The cell stocks used in the present studies were donated to the Kotton laboratory for the purpose of providing reference data. Samples were isolated by the overnight culture of lung explants in Waymouth media; a technique that generally yields 86 ± 2% epithelial cells with the remaining cells consisting of fibroblasts with <1% endothelial cells.

To isolate human primary lung epithelial cells, 1×1cm pieces of distal human lung obtained from healthy regions of the upper lobe of non-utilized human lungs donated for transplantation were dissected and all airway tissue and pleura was resected. Tissue was digested using dispase, collagenase I, and DNase using the gentle MACS 63 dissociator (Miltenyi) for 30 minutes at 37°C. The cell suspension was passed over 70uM and 40uM filters to generate a single cell suspension. Magnetic bead sorting using MACS LS columns (Miltenyi) and the following antibodies: HTII-280 (anti-human AEC2 antibody, IgM, Terrace Biotechnologies) and anti-IgM magnetic beads (Miltenyi) was used to obtain purified human AEC2 cells and were subsequently collected into trizol.

#### Bulk RNA Sequencing

Sequencing libraries were prepared from the total RNA extracts of the above 13 samples using Illumina TruSeq RNA Sample Preparation Kit v2. The mRNA was isolated using magnetic beads-based poly(A) selection, fragmented, and randomly primed for reverse transcription, followed by second-strand synthesis to create double-stranded cDNA fragments. These cDNA fragments were then end-repaired, added with a single ‘A’ base, and ligated to Illumina® 64 Paired-End sequencing adapters. The products were purified and PCR-amplified to create the final cDNA library. The libraries from individual samples were pooled in groups of four for cluster generation on the Illumina cBot using Illumina TruSeq Paired-End Cluster Kit. Each group of samples was sequenced on each lane on the Illumina HiSeq 2500 to generate more than 30 million single end 100-bp reads.

Fastq files were assessed for quality control using the FastQC program. Fastq files were aligned against the human reference genome (hg19/hGRC37) using the STAR aligner (Dobin et al., 2013). Duplicate reads were flagged using the MarkDuplicates program from Picard tools. Gene counts represented as counts per million (CPM) were computed for Ensembl (v67) gene annotations using the Rsubread R package with duplicate reads removed. Genes with 10% of samples having a CPM < 1 were removed and deemed low expressed. The resultant data was transformed using the VOOM method implemented in limma R package (Law et al., 2014). Voom transformed data was then tested for differential gene expression using standard linear models using the limma package. Multiple hypothesis test correction was performed using the Benjamini–Hochberg procedure (FDR). Heatmaps and PCA plots were generated in R. All raw fastq files are available on-line at GEO (GSE131768).

### Directed differentiation of type 2 alveolar cells from PSCs

#### iPSC Line Generation and Maintenance

All experiments involving the differentiation of human PSC lines were performed with the approval of the Institutional Review Board of Boston University (protocol H33122). The BU3 iPSC line carrying NKX2-1^GFP^ and SFTPC^tdTomato^ reporters (BU3 NGST) was obtained from our prior studies (Hawkins et al., 2017; Jacob et al., 2017). This line was derived from a normal donor (BU3) (Kurmann et al., 2015). All PSC lines used in this study (BU3, RUES2, SPC2, and ABCA35) displayed a normal karyotype (BU3, 46XY; RUES2, 46XX; SPC2 46XY; and ABCA35 46XX) when analyzed by G-banding both before and after gene-editing (Cell Line Genetics). The human embryonic stem cell line RUES2 was a generous gift from Dr. Ali H. Brivanlou of The Rockefeller University. All PSC lines were maintained in feeder-free conditions, on growth factor reduced Matrigel (Corning) in 6-well tissue culture dishes (Corning), in mTeSR1 medium (StemCell Technologies) using gentle cell dissociation reagent for passaging. Further details of iPSC derivation, characterization, and culture are available for free download at http://www.bu.edu/dbin/stemcells/protocols.php.

#### Directed Differentiation of PSCs

As previously described (Jacob et al., 2017) we performed PSC directed differentiation via definitive endoderm into NKX2-1 lung progenitors as follows. In short, cells maintained on mTESR1 media were differentiated into definitive endoderm using the STEMdiff Definitive Endoderm Kit (StemCell Technologies) and after the endoderm-induction stage, cells were dissociated with gentle cell dissociation reagent (GCDR) and passaged into 6 well plates pre-coated with growth factor reduced Matrigel in ‘‘DS/SB’’ anteriorization media, consisting of complete serum-free differentiation medium (cSFDM) base as previously described (Jacob et al., 2017) supplemented with 10 µm SB431542 (‘‘SB’’; Tocris) and 2 µm Dorsomorphin (‘‘DS’’; Stemgent). For the first 24 hr after passaging, 10 µm Y-27632 was added to the media. After anteriorization in DS/SB media for 3 days (72 hr), cells were cultured in ‘‘CBRa’’ lung progenitor-induction media for 9-11 days. ‘‘CBRa’’ media consists of cSFDM containing 3 µm CHIR99021 (Tocris), 10 ng/mL recombinant human BMP4 (rhBMP4, R&D Systems), and 100nM retinoic acid (RA, Sigma), as previously described (Jacob et al., 2017). On day 15 of differentiation, live cells were sorted on a high-speed cell sorted (MoFlo Legacy or MoFlo Astrios EQ) based on GFP expression for further differentiation or analysis as indicated in the text.

Sorted day 15 or 17 cells (as described in the text) were resuspended in undiluted growth factor-reduced Matrigel (Corning) at a dilution of 500 cells/µL, with droplets ranging in size from 25 to 100µL in 12 well tissue culture-treated plates (Corning). Cells in 3D Matrigel suspension were incubated at 37°C for 20-30 min, then warm media was added to the plates. Where indicated in the text, outgrowth and distal/alveolar differentiation of cells after day 15 was performed in ‘‘CK+DCI+Y’’ medium, consisting of cSFDM base, with 3 µm CHIR99021, 10 ng/mL rhKGF, and 50 nM dexamethasone (Sigma), 0.1mM8-Bromoadenosine 30,50-cyclic monophosphate sodium salt (Sigma) and 0.1mM3-Isobutyl-1-methylxanthine (IBMX; Sigma) (DCI) and 10 µm Y-27632. For NanoString mRNA analysis, alveolospheres were released from Matrigel droplets, and for flow cytometry and cell sorting, they were dissociated into single cell suspension. To release alveolospheres from Matrigel, droplets were incubated in dispase (2mg/ml, Fisher) at 37°C for 1 hr, centrifuged at 300 g x 1 min, washed in 1x PBS, then centrifuged again at 300 g x 1 min. To generate single cell suspensions, cell pellets were incubated in 0.05% trypsin and continued through the trypsin-based dissociation protocol described above, after which they could be passaged into fresh Matrigel and analyzed by flow cytometry.

#### NanoString Time Series

We used NanoString nCounter for direct quantification of 66 genes in triplicate of PSC differentiations based on a high-frequency sampling design: day 0, 17, 19, 21, 23, 25, 27, 29, 31, 33 with n=3 late human fetal lung alveolar epithelial cell samples included as controls (Late HFL; 21 weeks gestation). RNA extraction was performed by miRNeasy MicroKit (Qiagen) following the manufacturer’s protocol. RNA concentration and integrity were measured using NanoDrop ND-2000 and 2200 Tape Station. The NanoString nCounter™ XT CodeSet Gene Expression Assay was performed using 100 ng total RNA as previously described (Herazo-Maya et al., 2017). The raw data was background corrected and normalized using NanoStringQCPro (Nickles et al., 2018). The estimation of non-specific noise for background correction was done using the signals obtained from negative controls in each lane. For content normalization we scaled each probe relative to the average signal from pre-annotated housekeeping genes. The average and range of the normalized values are shown in Figure 2 and Figure S1.

### Time Point Selection (TPS) Analysis

We used the Time Points Selection (TPS) method(Kleyman et al., 2017) to determine time points to profile for accurate model reconstruction. Briefly, TPS utilizes NanoString nCounter quantification to obtain a densely sampled subset of genes which are known to be relevant to the process (in this case, known lung development genes; see figures 2 and supplement). It then uses a greedy algorithm to identify the best time points to use given a limited budget (i.e. if the user can only profile x number of time points). TPS can also estimate the resulting error from using less time points and so provides a way to balance accuracy and costs. Here we searched for different values of x ranging from 4 to 8. As Figure 2 shows, we observe an elbow in the error plot when using 6 time points. Such elbow means that further increase in the number of time points does not lead to much decrease in error. For 6 time points the expected error (0.21, log2 difference) is not far from the expected error due to repeats (0.16) which is the optimal error we can obtain and likely reflects real biological variations or technical issues. We thus used the 6 selected time points (15, 17, 21, 25, 29 and 31) to profile the single cells.

### Time Series Single-Cell Transcriptomic Analysis of AEC2 Directed Differentiation

#### Time series cell capture and profiling by scRNA-seq with SPRING plot visualization

Day 15 BU NGST cells were sorted for NKX2-1^GFP^ (as described above) and live positive and negative sorted cells were acquired for scRNA-seq in the Harvard Medical School (HMS) scRNA-seq core laboratory. NKX2-1^GFP^ positive cells were replated in 3D matrigel and grown for a further 16 days to day 35. At the selected time points (days 15, 17, 21, 25, 29 and 31) cells derived from directed differentiation of the BU3 NGST iPSC line were stained with calcein blue viability dye and sorted to obtain live cells for scRNA-seq as detailed in the text. Cells were captured in the Harvard Medical School (HMS) Single Cell Core for scRNA-seq using inDrops technology, as follows. First cells were assessed for cell number and viability and resuspended in 1000 µL of 15% OptiPrep™ to allow for a homogenous resuspension and reduced clumping. Cell capture and library preparation were performed using a modified version of inDrops protocols (Klein et al., 2015; Plasschaert et al., 2018) involving encapsulation of cells into 3-nl droplets with hydrogel beads carrying barcoding reverse transcription primers. Following the within-droplet reverse transcription step, emulsions were split into batches of approximately 2,000 cells, frozen at −80C, and subsequently processed as individual RNA-seq libraries (see Table S10). Approximately 4,000 cells for each time point were encapsulated for scRNA-seq.

The standard transcriptome RNA-seq libraries were processed as previously reported (R.Zilionis et al., 2017). In brief, the single cell libraries were demultiplexed following the recommended inDrops pipeline (https://github.com/indrops/indrops) in order to generate count matrices for each sample. We used the repeat-masked primary assembly of the human genome GRCh38 (ENSEMBL) as a reference. Reads were filtered according to the protocol to remove those that had low quality or low complexity. After counting and sorting abundant barcodes, histograms were used to identify thresholds that separate cells from empty gel beads. Finally, the reads of each barcode were aligned to the reference genome with Bowtie. Next, demultiplexed count matrices of the four libraries were aggregated into one combined analysis for downstream analysis. After further filtering to remove putative doublets as well as stressed or dying cells (having >20% or UMIs coming from mitochondrial genes), we performed linear dimensionality reduction with PCA, which was then used as input for Louvain clustering and non-linear dimensionality reduction with tSNE. Cell cycle stage was scored and classified using the strategy described in (Tirosh et al., 2016). Differential expression was tested using hurdle models for sparse single-cell expression data implemented in MAST (Finak et al., 2015). The derived markers were used to annotate the identity of each cluster. Clonal identity was derived from the lineage barcoding spiked samples. The association between cellular barcodes and lentiviral barcodes from the spiked samples was connected to the transcriptomic samples for visualization. Lineage-annotated transcriptome data was then imported into SPRING (Weinreb et al., 2018a) for interactive analysis and visualization. All SPRING plots (k-NN graphs rendered using a force-directed layout) were generated in the SPRING upload server:https://kleintools.hms.harvard.edu/tools/spring.html using the default parameters: 0 minimum UMI total for filtering cells, minimum of 3 cells with >= 3 counts for filtering genes, 80 percentile as threshold of gene variability for filtering genes, 50 PCA dimensions for building graph, and a k of 5 for the k-Nearest Neighbours algorithm used to create the graph. Annotation tracks (clusters) were imported from upstream analysis with the Seurat package (Louvain algorithm at resolution 0.6) and gene sets for predicting the degree of maturation (6 gene set) and differentiation (8 gene set) were derived from the bulk RNA-seq data analysis (see Figure 1). The final figures were plotted using ggplot2 package (R-CRAN) and edges between nodes omitted for clarity. Datasets are available for download from GEO (GSE137811).

### Predicting and mapping fate trajectories using Continuous State Hidden Markov Models (CSHMM)

We used Continuous State Hidden Markov Models (Lin and Bar-Joseph, 2019) to reconstruct the branching process of the data. The model was first initialized by clustering cells at each time point. Next, the location of each cluster was adjusted based on the distance to the root cluster (D15) to account for asynchronous development among cells at the same time point. In our case this led to placing cells from Day 21 and 25 at the same level of the branching tree because most overlapped in terms of expression (See also Spring plot in Figure 3 and Figure S2). For similar reasons we grouped cells at D29 and D31 at the same level. After assigning initial level, clusters in each level were connected to the nearest cluster in the previous level where distance is based on expression similarity leading to a tree like branching structure. Finally, for the initializations, cells in each cluster were randomly placed on the paths connecting their node to their parent node (Figure 4A).

Next, CSHMM learned parameter values and cell assignments using an Expectation Maximization (EM) algorithm. In our CSHMM setting, neighboring states along a path share parameters and so, while the total number of states is potentially infinite the number of model parameters is still finite. The model is further constrained by allowing transition to only a finite (though not necessarily small) number of states from each state as shown in Schematic below. Parameters (learned in the M step) include standard HMM parameters as were used in previous bulk modeling methods (Ernst et al., 2007) and a new, gene specific parameter vector denoted by Kx (where x is the path number) which allows genes to change differently along a path. The structure and cell assignments also provide emission probabilities for each state. This enables the method to assign gene specific expression profiles for each of the paths while cell assignments, determined in the E step, are continuous and so cells can be assigned to any point along the path. Given the constraints on several aspects of the model, learning and inference is still efficient despite the infinite many possible states in a CSHMM. We stop when the likelihood of the model does not increase (Figure 4A). See (https://www.biorxiv.org/content/biorxiv/early/2018/07/30/380568.full.pdf) for complete details.

### Schematic: CSHMM model structure and parameters

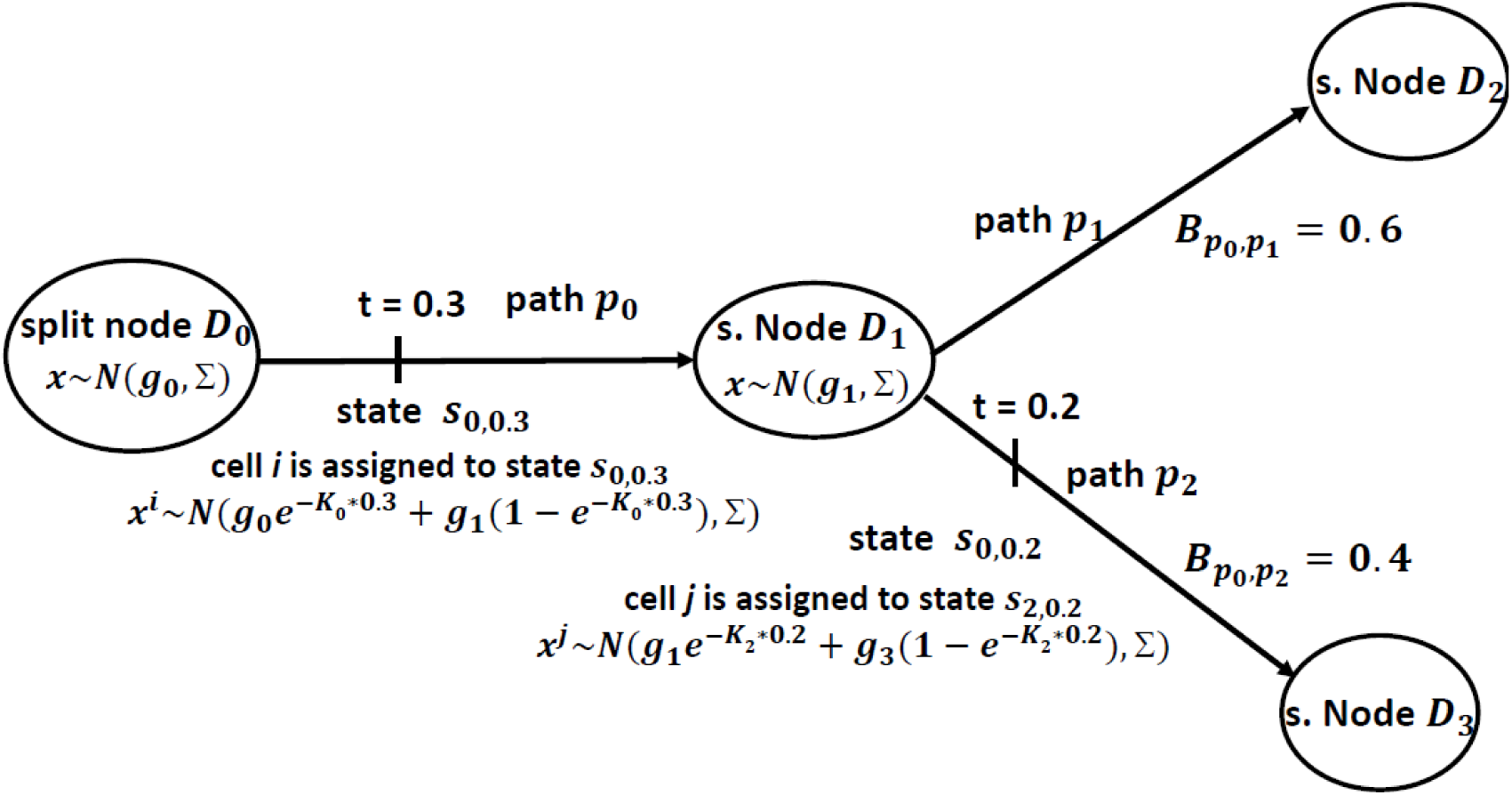

Each path represents a set of infinite states parameterized by the path number and the location along the path. For each such state we define an emission probability and a transition probability to all other states in the model. Emission probability for a cell along a path is a function of the location of the state and a gene specific parameter for each gene in the cell which controls the rate of change of its expression along the path. Split nodes are locations where paths split and are associated with a branch probability. Each cell is assigned to a state in the model. Further details available in: https://www.biorxiv.org/content/biorxiv/early/2018/07/30/380568.full.pdf

### CSHMM Prediction of Optimal Wnt Withdrawal Time and Confirmation by Direct Experiment

#### Determining time of split between cell fates

We used CSHMM assignments to infer the most appropriate time to withdraw Wnt. For this we selected a number of Wnt markers detailed in the text and plotted their continuous expression along the top and bottom paths (**Error! Reference source not found.**. We used these plots to determine an accurate split time, in the model, for these markers. To determine the actual time the model point corresponds to the trajectory split, we assigned each *N* node in the CSHMM a real time which is computed by averaging the time in which cells assigned right before and right after the node were profiled (Figure 5B). Using these values we assigned time to each point along a path by interpolating the time assigned to the two N nodes that define the path. For example, the split point identified for Wnt is in the middle of the N1-N2 path. Since N1 is assigned to day 16 and D2 to day 19, this point is assigned to day 17.5.

#### Direct Wnt Withdrawal During NKX2-1+ Outgrowth Differentiation

Day 15 BU NGST cells were sorted for NKX2-1^GFP^ (as described above) and live positive sorted cells were replated in 3D Matrigel and grown for a further 14 days to day 29. We withdrew the GSK antagonist, CHIR from our “CK+DCI” media for a period of 4 days, starting at five different time points (days 17,19, 21, 23 and 25; Figure 5C). To maintain proliferation of resulting cells CHIR was added back after 4 days, allowing each parallel condition to be harvested at the identical total differentiation time while keeping the length of CHIR withdrawal (4 days) constant for each condition. At day 29 a single cell suspension of all alveolospheres was generated and cells were analyzed by flow cytometry for NKX2-1^GFP^ and SFTPC^tdTomato^ reporter expression.

#### Knockdown of CEBPδ by siRNA transfection

The iPSC line, SPC2 was differentiated to alveolospheres as described above. Spheres were dissociated to single cells with trypsin, washed and counted. Cells (5×10^5^ cells per reaction) were resuspended in a 20 uL nucleofection reaction, as per manufacturer’s instructions (Lonza) (16.4 uL P3 solution with 3.6 uL supplement), with 500 nM of non-targeting siRNA (Dharmacon, #D-001810-01-05) or CEBPδ siRNA (Dharmacon, #L-010453-00-0005). Cells were transferred to a cuvette and nucleofected with program EA104 (4D Nucleofector; Lonza). Each reaction was replated in 100 uL 3D Matrigel in CK+DCI+Y. Cells were collected, by dissolving Matrigel with dispase, at 48 hours for RNA isolation. CEBPδ (Hs00270931_s1), SFTPC (Hs00161628_m1), SFTPB (Hs00167036_m1), ABCA3 (Hs00184543_m1), SFTPA1 (Hs01652580_g1), NKX2-1 (Hs00968940_m1) and FOXA2 (Hs00232764_m1) transcripts were quantified by qRT-PCR and fold-change was calculated with respect to the non-targeting siRNA group as described in the section below.

### Lineage Tracing of PSC-derived iAEC2 Differentiation Using Lentiviral Barcoding

#### Lentiviral barcoding

Lineage tracing of individual cells was performed using a lentiviral barcode labeling system(Weinreb et al., 2019). Human BU3 NGST cells were differentiated, sorted on D15 for NKX2-1^GFP^ positive cells and grown as alveolospheres, as described above, until D17. On D17 the Matrigel matrix was dissociated using 2mg/ml Dispase solution (Thermofisher, 17105041) and 6.5e4 cells were resuspended in 600ul CK+DCI+RI with polybrene (5ug/ml). This cell suspension was divided equally into two suspensions: an uninfected control (MOI=0) and a sample containing 32,500 cells which was infected with lentivirus (MOI=10). Both samples were left in suspension for 4 hours before being washed and replated. Each sample was resuspended as cell clumps in 50ul Matrigel (Corning 356231) and plated as two 25 ul droplets in one well of a 12-well plate previously coated with 100ul of Matrigel (Corning 356231). Cells were fed with CK+DCI+RI every 2 days until collection on D27, at which point cells were collected for single cell RNA-sequencing. Cells were collected using a MoFlo Astrios EQ cell sorter and enriched using calcein blue, to sort out live cells, and/or viral GFP expression.

Cell capture and library preparation were performed as described using a modified version of inDrops protocols(Klein et al., 2015; Plasschaert et al., 2018). Prior to library preparation, RNA fractions generated from each population were split in half, with one half being used for standard library prep and the other half for targeted lineage barcode enrichment. To enrich for lineage barcodes, library preparation was modified as previously described (Weinreb et al., 2019) by skipping RNA fragmentation, priming the RT reaction using a barcode-specific primer (TGAGCAAAGACCCCAACGAG), introducing an extra PCR step using a targeted primer (8 cycles using Phusion 2X master mix; Thermofisher; primer sequence = TCG TCG GCA GCG TCA GAT GTG TAT AAG AGA CAG NNN Ntaa ccg ttg cta gga gag acc atat), and 1.2X bead purification (Agencourt AMPure XP). All targeted and non-targeted final libraries were pooled at equimolar ratios and sequenced using Illumina NextSeq 500 Sequencing (75 Cycles, 75bp Single Read sequencing).

The lentivirus-targeted single-cell libraries were demultiplexed in the same way as the transcriptome libraries and further processed following the LARRY pipeline (Lineage And RNA RecoverY) described in (Weinreb et al., 2019). Briefly, this involves two steps: the first step was sorting and filtering the raw sequencing reads generated from the inDrops pipeline (https://github.com/indrops/indrops) which provides a list of reads with annotated cell barcode and unique molecular identifier (UMI); we used, as a threshold for collapsing lentivirus-barcodes, a hamming distance of 3, and filtered out cell-lineage combinations that were not supported by at least 10 reads. The second step was annotating the clonality of cells, with further stringent filtering to discard contaminated droplets, which resulted in a NxM binary matrix of 0/1, where N is number of cells and M is the number of clones. The pipeline was executed using the implementation developed by Klein Lab, available online at: https://github.com/AllonKleinLab/LARRY. Datasets are available for download from GEO (GSE137805).

#### Projection of Barcoded Cells on CSHMM

Given experimental differences in profiling cells used to construct the model and barcoded cells we compared bar-coded cell expression values using a subset of 82 genes. These included the top 68 DE genes between the top and bottom paths (P2 and P3) and 14 known lung cell markers (Table S4). In the CSHMM each location along a path is defined by an emission probability and so for each location we estimated the average expression value for each of these 82 genes. To assign bar-coded cells to the model we compared the expression of these genes to a densely sampled set of locations on each path (100 uniform locations). We assigned cells to the location which minimized the Euclidian distance between the bar-coded gene expression and the average expression learned for that location.

To determine the accuracy of our assignments we performed statistical tests based on randomization in which we sample a random expression profile and attempt to assign it to one of the paths in the same way we assigned the bar-coded cells. Using both t-test and ranksum test on the similarity of the projected profiles to the locations they were assigned to, we concluded that bar-coded cells are significantly associated with the paths they are assigned to, further supporting our general conclusion regarding late cell fate commitment.

### Determining Fate Retention in Additional iPSC lines

For independent validation of stable SFTPC^tdTomato+^ outgrowth by scRNA-seq using iPSCs from a variety of genetic backgrounds, two additional iPSC lines, “SPC2” and “ABCA35” (clones SPC2-ST-B2 and ABCA3_W308R ST13CR17Corr18), were obtained from the Boston University CReM’s iPSC Core Facility (Boston, MA). These lines were generated by reprogramming patient-derived fibroblasts (SPC2-18 and ABCA35 from Washington University; generous gift of Drs. F. Sessions Cole, Aaron Hamvas, and Jennifer Wambach, St. Louis, MO). The Institutional Review Board of Washington University, St. Louis, MO, approved procurement of these fibroblasts with documented informed consent. SPC2-18 cells were reprogrammed using the excisable, floxed lentiviral STEMCCA vector, with successful STEMCCA excision confirmed prior to directed differentiation as previously published (Somers et al., 2010). ABCA35 cells were reprogrammed with the Sendai reprogramming system (CytoTune, Thermofisher, Grand Island, NY). SPC2 cells originally carried a SFTPC^I73T^ heterozygous mutation and ABCA35 cells originally carried ABCA3^W308R^ homozygous mutations. After reprogramming, both lines underwent CRISPR gene editing to correct these mutations to generate control iPSC lines (K. Alysandratos et al. and Y. Sun et al. manuscripts in preparation). For tracking distal lung differentiation efficiency each line was engineered to carry a tdTomato reporter targeted to the endogenous SFTPC locus (SFTPC^tdTomato^ or “ST”) using previously published methods (Jacob et al., 2017).

### Isolation of SFTPC^tdTomato^-expressing iAEC2s and Long-Term Culture of Alveolospheres

To test for long term maintenance of the lung epithelial program in iPSC-derived alveolospheres we used our published iAEC2 differentiation protocol (Jacob et al., 2017) with extensive details including serial alveolosphere passaging techniques detailed in Jacob et al. 2019, Nature Protocols, in press. In brief, SPC2 iPSCs were differentiated until day 16 when primordial lung progenitors were sorted based on CD47^hi^/CD26^lo^ gating (Hawkins et al., 2017). After replating these purified progenitors in 3D Matrigel cultures in “CK+DCI” media, the resulting epithelial spheres were passaged without further sorting on day 37 and CHIR was briefly withdrawn from days 40-44 to achieve iAEC2 maturation (Jacob et al., 2017). Then CHIR was added back for the duration of the experiment. On day 51 SFTPC^tdTomato+^ cells were sorted and replated as alveolospheres with subsequent passaging *without further cell sorting* performed on days 65, 82, 101, and 115. scRNA-seq of all calcein blue-stained live cells was performed on day 115 using the 10X Chromium system with v2 chemistry as previously published (McCauley et al., 2018). Library preparation, sequencing, alignment and analyses were performed as previously published and tSNE plots with Louvain clustering and identity gene overlays prepared using our previously published pipeline (McCauley et al., 2018). Datasets are available for download from GEO (GSE137799).

### Quantification and Statistical Analysis

#### Reverse Transcriptase Quantitative PCR

RT-qPCR was performed as previously described (Hawkins et al., 2017). Briefly, RNA was isolated according to manufacturer’s instructions using the QIAGEN miRNeasy mini kit (QIAGEN). cDNA was generated by reverse transcription of up to 150ng RNA from each sample using the Applied Biosystems High-Capacity cDNA Reverse Transcription Kit. For qPCR, technical triplicates of each of at least 3 biological replicates were run for 40 cycles as either 20 uL reactions (for use in Applied Biosystems StepOne 96-well System) or 12 uL reactions (for use in Applied Biosystems QuantStudio7 384-well System). All primers were TaqMan probes from Applied Biosystems (see all in Key Resources Table). Relative gene expression was calculated based on the average cycle (Ct) value of the technical triplicates, normalized to 18S control, and reported as fold change (2(-DDCT)), with a fold change of 1 being assigned to untreated cells depending on the experimental conditions. Samples with undetectable expression after 40 cycles were assigned a Ct value of 40 to allow for fold change calculations.

#### Statistical Methods

Statistical methods relevant to each figure are outlined in the figure legend. In short, unless indicated otherwise in the figure legend, unpaired, two-tailed Student’s t tests were used to compare quantitative analyses comprising two groups of n = 3 or more samples, where each replicate (‘‘n’’) represents either entirely separate differentiations from the pluripotent stem cell stage or replicates differentiated simultaneously and sorted into separate wells. Further specifics about the replicates used in each experiment are available in the figure legends. In these cases, a Gaussian distribution and equal variance between samples was assumed as the experiments represent random samples of the measured variable. The p value threshold to determine significance was set at p = 0.05. Data for quantitative experiments is typically represented as the mean with error bars representing the standard deviation or standard error of the mean, depending on the experimental approach. These details are available in the figure legends.

## DATA AND SOFTWARE AVAILABILITY

The data discussed in this publication have been deposited in NCBI’s Gene Expression Omnibus (Edgar et al., 2002) and are accessible through GEO Series accession numbers GSE131768, GSE137799, GSE137805 and GSE137811, and is also available on the Kotton Lab’s Bioinformatics Portal at http://www.kottonlab.com. Software for CSHMM is available at: https://github.com/jessica1338/CSHMM-for-time-series-scRNA-Seq and an interactive webtool for the CSHMM of the time series data is publically accessible online at: cosimo.junding.me

## ADDITIONAL RESOURCES

Further protocol information for reprogramming, iPSC/ESC cultures, directed differentiation and the production of lentiviral particles can be found at http://www.bu.edu/dbin/stemcells/protocols.php.

## Supplemental Information titles and legends

**Table S1.** Top 1000 most varying genes in primary developing human AEC2s from fetal to adult (weeks 16 of gestation to adult) (see downloadable excel sheet): The list represents the top 1000 most varying genes with the last column representing gene variance. The log FC column in the expression spreadsheet indicates the log2 fold change compared to all other samples.

**Table S2.** List of top up-regulated genes for each of the paths (P0-P10) in the CSHMM (see downloadable excel sheet):

In all 11 sub-tables the up-regulated genes in the cells of a specific path compared with all other cells are shown. To define up-regulated genes, we required log2 fold change >0.6 (∼1.5x) and p-value>0.05. In each of the sub-tables, the first column (gene) represents the up-regulated genes; the second column (Other_cells) represents the expression in all other cells; the third column (P0_Cells) shows the expression in the cells of the specific path; the last column (log2_fold_change) represents the log2 fold change. All expression is in log2 space.

**Table S3. Up-regulated genes in bottom paths (P3,P4,P6,P9) versus top paths (P2,P5,P7,P8,P10) (see downloadable excel sheet):** All the columns here have similar meanings as in Table S2. The first column (gene) represents the up-regulated genes in bottom paths; the second column (Top_paths) represents the expression in Top paths; the third column (Bottom Paths) represents the expression in Bottom paths; the last column (log2_fold change) denotes the log2 fold change of gene expression between top paths and bottom paths.

**Table S4.** List of top DE genes for each clusters for all cells profiled in the lentiviral barcoding scRNA-seq lineage tracing experiment (see downloadable excel sheet): **Top** 20 differentially expressed genes (FDR < 0.05, ranked by log2 fold-change) for each cluster from the lineage tracing experiment. Column pct.1 and pct.2 refer to percentage of cells expressing each gene in the cluster of interest and other respectively.

**Table S5.** Lentiviral barcoded clone cells mapped to bottom (P3,P4,P6,P9) Versus top paths (P2,P5,P7,P8,P10) (see downloadable excel sheet): The first column (index) represent the Lentivirus clone Index; the second column (size) denotes the number of cells in the clone; the P0-P11 columns show the number of cells in each of the paths; the last column denotes ratio of cells mapped to bottom paths versus cells mapped to the top paths.

---

**Figure S1.**
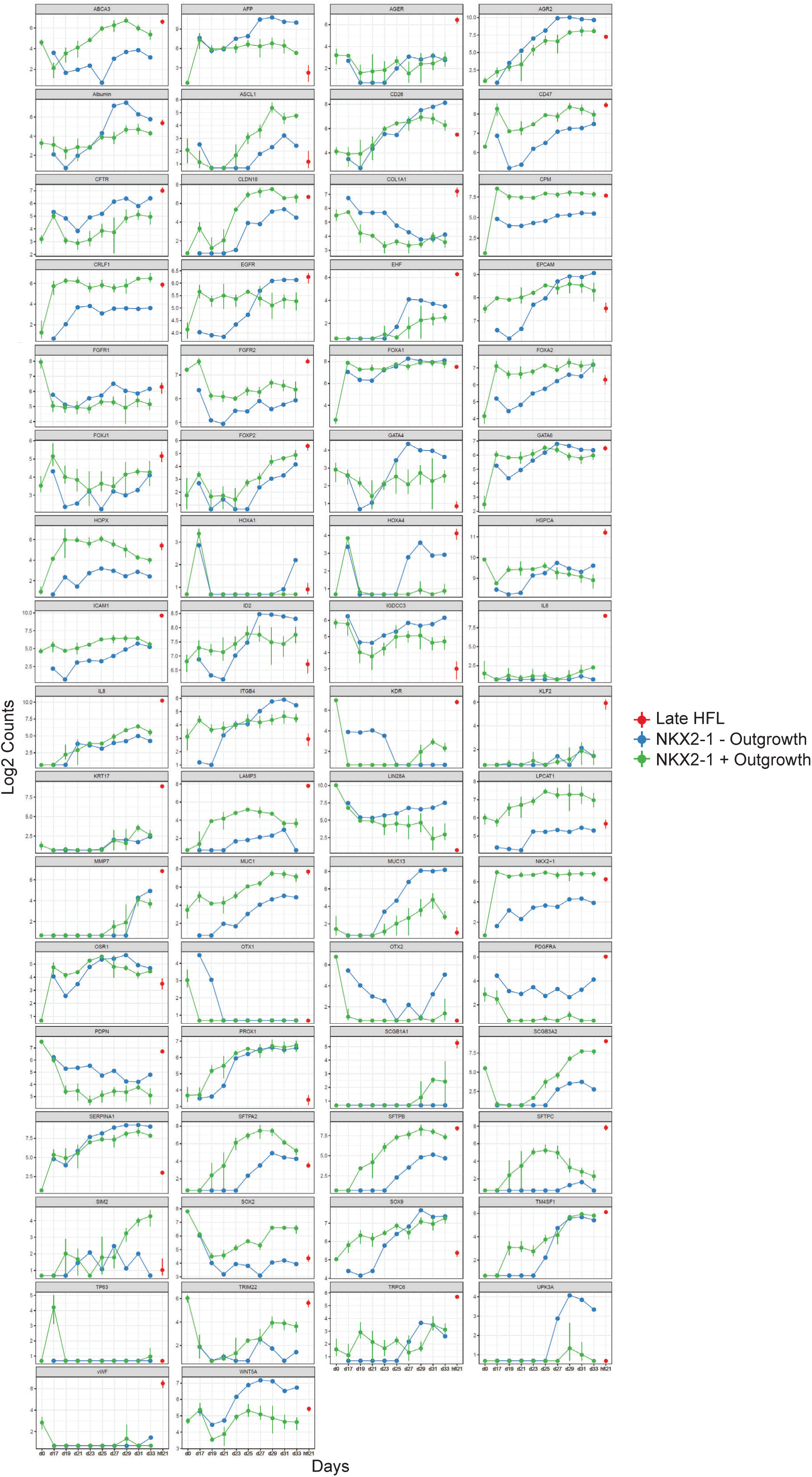
NanoString gene expression time series of iAEC2 differentiation. Average and range of expression over time for the entire 66 gene panel used to monitor AEC2 directed differentiation in vitro. The panel includes markers of AEC2 maturation/differentiation, proximal lung markers, non-lung endoderm markers, mesenchymal, and other non-lung markers for cells derived from PSCs (0 to 33 days of differentiation, n=3 biological replicates form distinct differentiation runs). Red dot indicates late human fetal lung controls (HFL; 21 weeks gestation).

**Figure S2.**
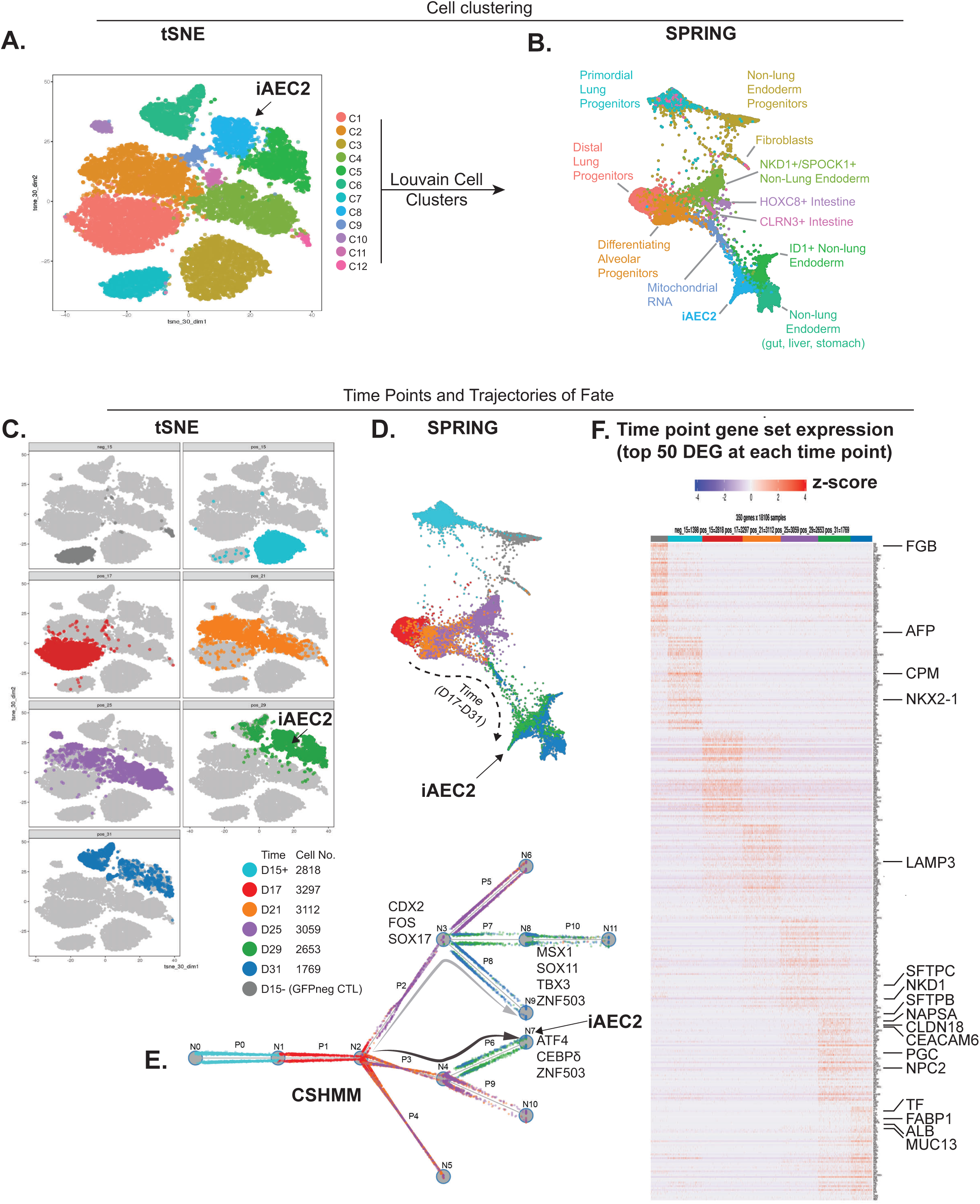
Fate trajectories of iPSC-directed differentiation are revealed by CSHMM. (A) tSNE representation of transcriptional profiles of cells from all scRNA-seq time points in the time series with 12 Louvain Clusters identified, including the blue annotated cluster ‘iAEC2’ containing multiple gene markers of AEC2 cells. (B) SPRING plot representing the transcriptional profiles of cells from all time points in the scRNA-seq time series with the same 12 Louvain Clusters from (A), and putative cluster identities annotated based on markers for lung and non-lung endoderm lineages. (C) tSNE representation (as in panel A) of the transcriptional profiles of all cells from the time series with each time point highlighted using individual colors (Grey, Day 15 NKX2-1 negative sort or control, ‘D 15 -’), (Cyan, Day 15 NKX2-1^GFP^ positive sort, ‘Day 15 +‘), and the resulting outgrowth of the D15 GFP+ sorted cells shown in the indicated color at each of *6 time points*. (D) SPRING plot demonstrating cells from each time point highlighted in the same individual colors shown in (C). (E) CSHMM branching model with each cell individually colored by time point, using the same colors shown in previous panels (C) and (D). Note both SPRING and CSHMM suggest differentiating cell fate trajectories whereas tSNE does not. (F) Heatmap of top 50 differentially expressed genes at each time point (positive fold change compared to cells in all other clusters, FDR<0.05) highlighted with the same colors shown in panels (C), (D) and (E)) with selected genes for lung and non-lung endoderm highlighted on the right side of the panel.

**Figure S3.**
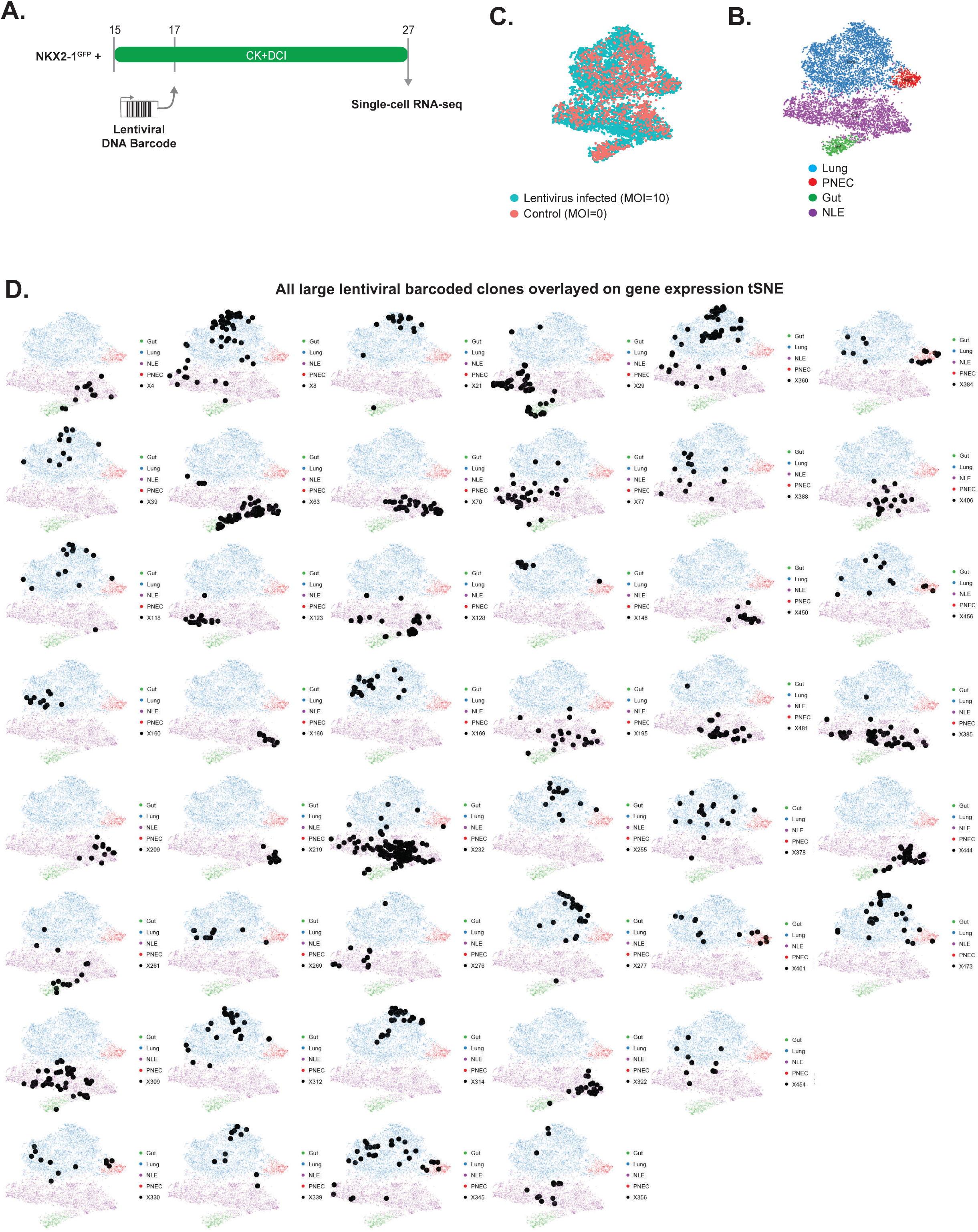
Lineage tracing of iPSC-derived type 2 alveolar cells with lentiviral barcoding and subsequent scRNA-seq. (A) Schematic of experiment showing infection of NKX2-1GFP+ sorted cells grown out to day 17 at which time cells are infected with a lentiviral library to tag individual cells with unique integrated DNA barcodes. After infection cells were replated in 3D Matrigel and grown for a further 10 days in CK+DCI media at which point a single cell suspension was generated and cells were encapsulated for scRNA-seq using inDrops. Inherited lentiviral barcodes were matched with transcriptomic profiles for each cell to track clones from day 17 to day 27. (B) tSNE plot of all cells harvested at day 27 with cells infected with lentivirus at multiplicity of infection (MOI) 10 highlighted in green and cells not infected (MOI 0) shown in orange, demonstrating that cells infected with lentivirus were found in all clusters. (C) tSNE plot of all cells harvested at day 27 with Louvain clustering (resolution 0.25) which has been annotated based on marker genes for distal lung alveolar epithelium ‘Lung’, pulmonary neuroendocrine cells (PNEC), Gut and non-lung endoderm (NLE). (D) tSNE plots with Louvain clustering for annotated cell lineage and overlayed with all 45 lentivirually tagged large clones (threshold for “large” is >10 cells per clone).

**Figure S4.**
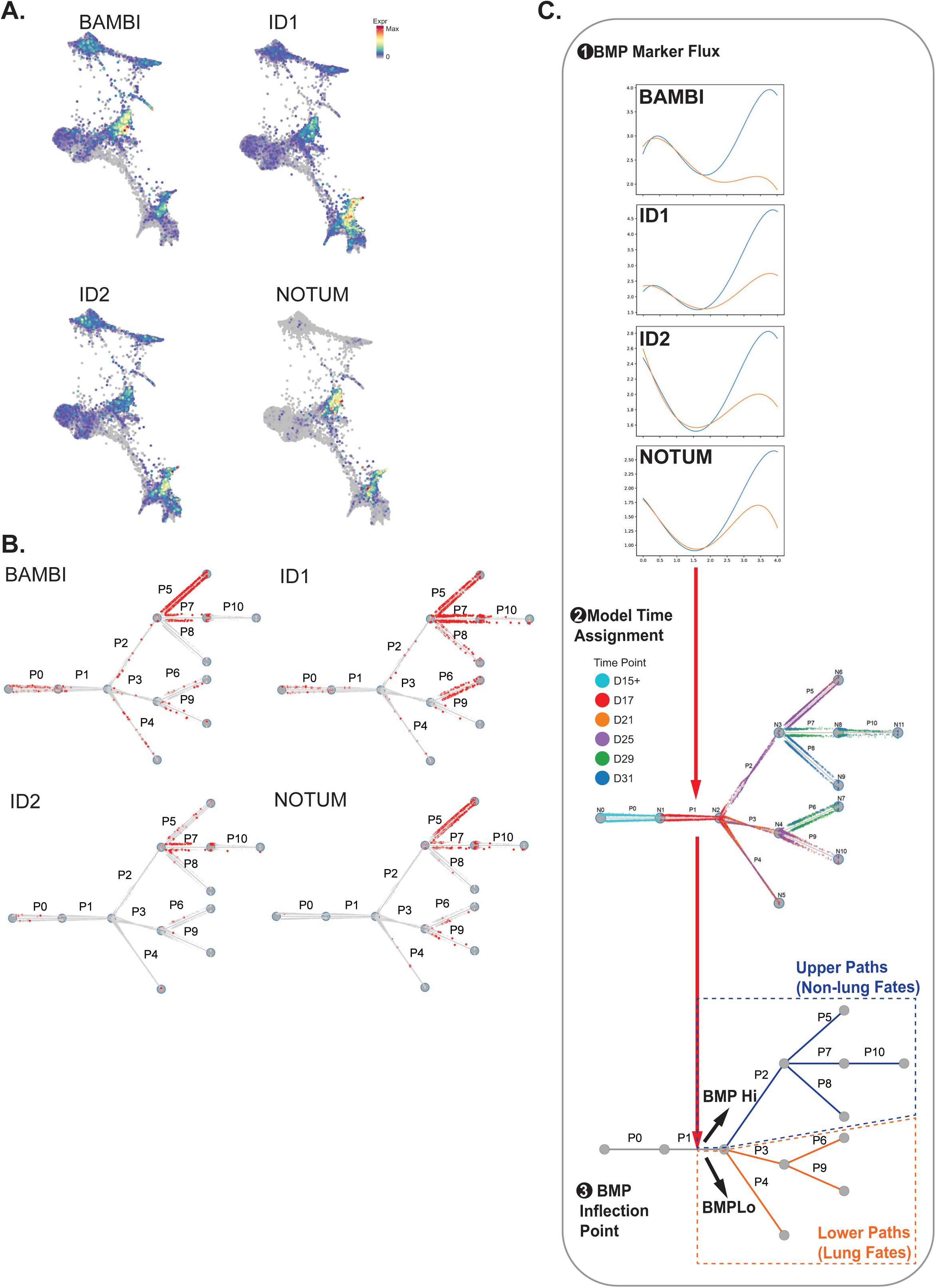
Increased BMP target gene expression in cells diverging to non-lung endodermal fates. (A) Normalized gene expression overlayed on SPRING plots for the key BMP target genes BAMBI, ID1, ID2 and NOTUM. (B) Expression of the same key BMP target genes in individual cells overlaid on CSHMM showing increased expression in upper paths (non-lung). (C) To determine the time of BMP pathway activation the continuous expression of these markers is reconstructed using splines to plot the reconstructed expression profiles for the four indicated BMP signaling markers for cells assigned to the top paths (blue curve) vs. bottom paths (orange curve). For all four there is a split in expression values with higher expression of BMP signaling markers present in those cells diverging to a non-lung fate trajectory (upper paths).

**Figure S5.**
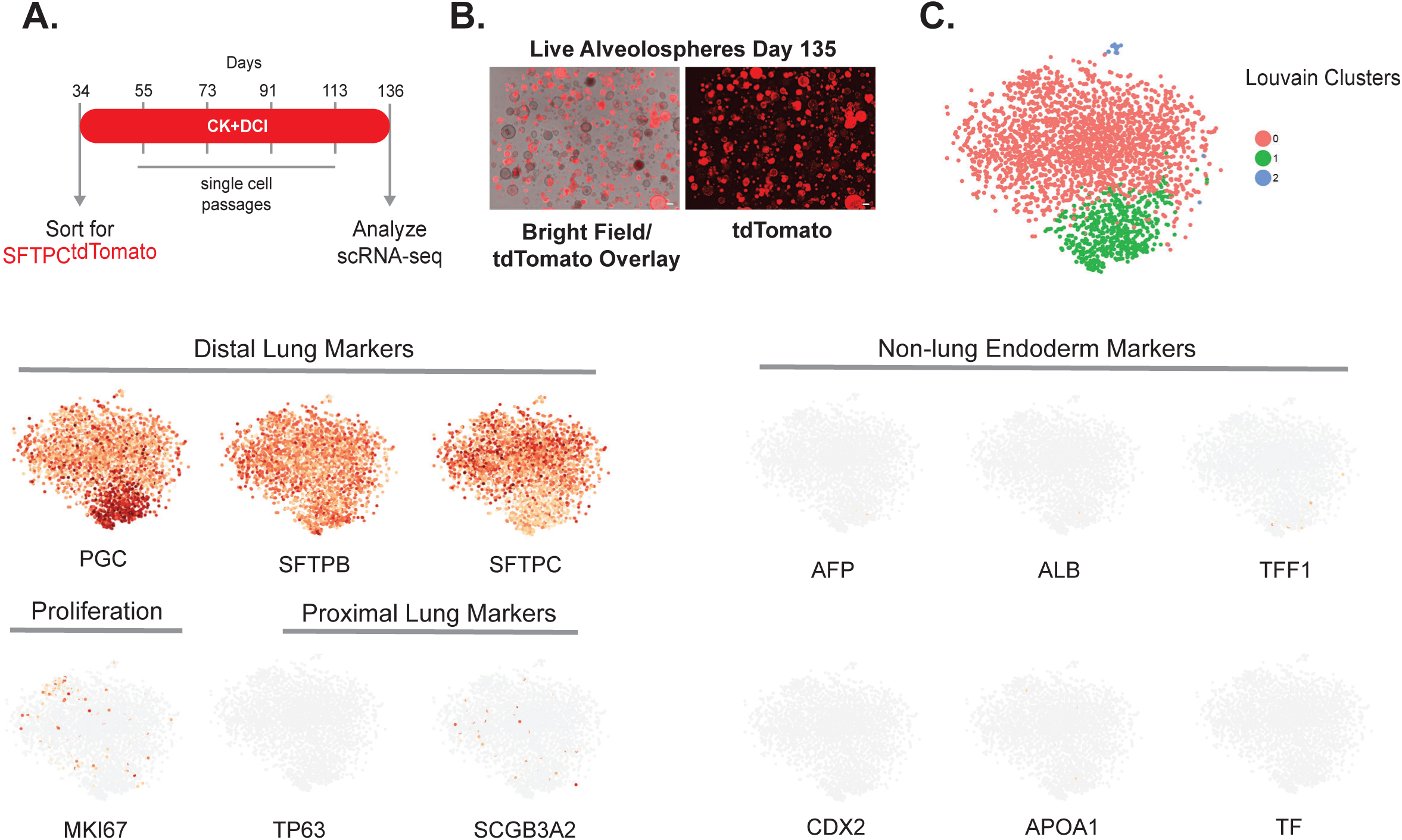
Time-dependent maturation is associated with retention of iAEC2 fate in the ABCA35 iPSC line. (A) Similar to Figure 7, pictured is a schematic of experiment in which cells sorted at day 34 for SFTPC^tdTomato^ were replated as alveolospheres in 3D conditions in CK+DCI media. They were subsequently passaged as single cells 4 times on the days indicated, without further cell sorting. At day 136 live cells were encapsulated for scRNA-seq using the 10X Chromium System. (B) Representative images of live ABCA35 alveolospheres (bright-field/tdTomato overlay (day 135) illustrating retention of lung fate, indicated by continued expression of SFTPC^tdTomato^. Scale bar, 300 µm. (F) tSNE with Louvain Clusters. (G) Normalized gene expression overlayed on tSNE plots for selected markers for ‘distal Lung’ (PGC, SFTPB, and SFTPC), ‘Proliferation’ (MKI67), ‘Proximal Lung’ (TP63 and SCGB3A2), or (H) ‘Non-Lung Endoderm’ (AFP, ALB, TFF1, CDX2, APOA1 and TF).

**Table S6.** Subset of genes used to assign lentiviral barcoded cells onto CSHMM. A subset of 82 genes, including the top 68 genes differentially expressed between the upper and lower paths (P2 and P3) and 14 distal lung markers from our primary cell datasets was used to assign lentiviral barcoded cells onto CSHMM. Genes and criteria for inclusion in the subset are listed in the table.

| **Gene Name** | **Criteria** |
| --- | --- |
| AHNAK | DE gene |
| ATP6V1G1 | DE gene |
| BAMBI | DE gene |
| CADM1 | DE gene |
| CDC42EP3 | DE gene |
| CDH1 | DE gene |
| CDH17 | DE gene |
| CDX2 | DE gene and marker gene |
| COL2A1 | DE gene |
| CRABP2 | DE gene |
| CST1 | DE gene |
| DPY19L1 | DE gene |
| DPYSL2 | DE gene |
| DSC2 | DE gene |
| DUSP6 | DE gene |
| ERRFI1 | DE gene |
| FAM84A | DE gene |
| FNDC3B | DE gene |
| FOS | DE gene |
| FOXA1 | DE gene |
| FTL | DE gene |
| GNAI1 | DE gene |
| HIPK2 | DE gene |
| HMGA2 | DE gene |
| HMGN2 | DE gene |
| HSPD1 | DE gene |
| ID1 | DE gene |
| IDS | DE gene |
| IGDCC3 | DE gene |
| IRS2 | DE gene |
| ITGA2 | DE gene |
| LAMC1 | DE gene |
| LGALS3 | DE gene |
| LIFR | DE gene |
| LIX1L | DE gene |
| MAN1A1 | DE gene |
| MLLT11 | DE gene |
| MSN | DE gene |
| MYH9 | DE gene |
| NEAT1 | DE gene |
| PHLDA2 | DE gene |
| PIP4K2C | DE gene |
| PITX1 | DE gene |
| PPIC | DE gene |
| PRTG | DE gene |
| PSAT1 | DE gene |
| PTGES3 | DE gene |
| PXDN | DE gene |
| RASSF3 | DE gene |
| RHOBTB3 | DE gene |
| ROR1 | DE gene |
| S100A10 | DE gene |
| SDC3 | DE gene |
| SDC4 | DE gene |
| SLC2A1 | DE gene |
| SORT1 | DE gene |
| SOX17 | DE gene |
| SPTBN1 | DE gene |
| STK17B | DE gene |
| TBX3 | DE gene |
| THBS1 | DE gene |
| TMSB10 | DE gene |
| TSPAN13 | DE gene |
| TTR | DE gene |
| TUBB2B | DE gene |
| UGCG | DE gene |
| WIF1 | DE gene |
| ZFP36L2 | DE gene |
| APOB | marker gene |
| BMP3 | marker gene |
| CPM | marker gene |
| FABP1 | marker gene |
| FGB | marker gene |
| LPCAT1 | marker gene |
| NAPSA | marker gene (parameter reconstructed) |
| NKX2-1 | marker gene |
| SFTPA1 | marker gene (parameter reconstructed) |
| SFTPA2 | marker gene (parameter reconstructed) |
| SFTPB | marker gene (parameter reconstructed) |
| SFTPC | marker gene (parameter reconstructed) |
| TF | marker gene (parameter reconstructed) |
| TFF1 | marker gene (parameter reconstructed) |

**Table S7.** Assignment of lentiviral barcoded cells to the CSHMM model. t-test and ranksum test for the significance of the assignment of lentiviral barcoded cells to the CSHMM model.

| Method\#random cells | 1000 | 2566 (#infected cells) | 10000 |
| --- | --- | --- | --- |
| t-test | 5.28e-71 | 6.19e-105 | 1.30e-124 |
| Ranksum test | 3.64e-71 | 1.85e-130 | 5.05e-204 |

**Table S8.** Agreement between supervised and unsupervised lenti-virus cell assignment. Proportion of lentiviral barcoded cells assigned to lung path / state by projection to the CSHMM model and by unsupervised dimensionality reduction, respectively. Results are presented for all cells in large lenti clusters (30+). As can be seen, the agreement between the two assignment methods is very good.

| Clone Size | Lenti cluster | CSHMM projection | Clonality analysis |
| --- | --- | --- | --- |
| 212 | X232 | 0.065326633 | 0.019323671 |
| 79 | X360 | 0.710526316 | 0.835443038 |
| 77 | X29 | 0.142857143 | 0.053333333 |
| 75 | X63 | 0.02739726 | 0 |
| 69 | X8 | 0.742424242 | 0.835820896 |
| 65 | X128 | 0.046153846 | 0 |
| 61 | X385 | 0.052631579 | 0.016666667 |
| 57 | X309 | 0.054545455 | 0.035714286 |
| 40 | X70 | 0.025 | 0 |
| 37 | X444 | 0.142857143 | 0 |
| 34 | X481 | 0.03030303 | 0.029411765 |
| 33 | X345 | 0.793103448 | 0.696969697 |
| 32 | X314 | 1 | 1 |
| 32 | X77 | 0.28125 | 0.129032258 |

